# The nitrite reductase activity of xanthine oxidoreductase sustains cardiovascular health

**DOI:** 10.1101/2024.11.01.621390

**Authors:** Nicki Dyson, Rayomand S. Khambata, Tipparat Parakaw, Gianmichele Massimo, Ngan HH. Khuat, Annika A. Noor, Lorna C. Gee, Ivy Lim, Umme Siddique, Andrew J. Sullivan, Jonathan W. Ho, Krishnaraj Rathod, Michael R. Barnes, Claudia P. Cabrera, Amrita Ahluwalia

**Author notes:** **To whom correspondence should be addressed:** Amrita Ahluwalia, Barts and The London Faculty of Medicine & Dentistry, Queen Mary University of London, Charterhouse Square, London EC1M 6BQ, United Kingdom. Contributed equally (Nicki Dyson, Rayomand S. Khambata).

## Abstract

**Background:** Xanthine oxidoreductase (XOR) is a multi-functional enzyme that metabolises purines generating uric acid and is a generator of reactive oxygen species. Both functions have been implicated in the pathogenesis of cardiovascular disease. More recently, a third function of XOR as a nitrite reductase has been identified and been shown to play a key role in the benefits of targeting the non-canonical pathway for nitric oxide (NO) generation in the cardiovascular disease setting. This effect has been specifically attributed to XOR dependent recovery of NO levels. However, whether XOR derived NO plays any role in maintaining cardiovascular homeostasis in health is unknown. To explore this, we used global and hepatocyte-specific *Xdh*-deleted mice to assess cardiovascular homeostasis.

**Methods:** *Xdh*^+/+^ and *Xdh*^+/-^, *Xdh^fl/fl^ and Xdh^fl/fl^AlbCre^+/-^*(HXOR KO) mice littermates, matched for sex and age, were used for in vivo cardiovascular phenotyping; blood pressure, cardiac function, endothelial reactivity, and leukocyte trafficking. Tissues from these mice were used for biochemical measurements of nitrate, nitrite, and markers of NO downstream signalling.

**Results:** *Xdh*^+/-^ and HXOR KO mice expressed significantly attenuated liver and plasma nitrite reductase activity and platelet cGMP levels versus littermate controls. These effects were associated with increased systolic blood pressure, left ventricular remodelling, and increased leukocyte activation. These effects were associated with and likely driven by endothelial dysfunction evident in both mouse models. This dysfunction was reflected by increased endothelial adhesion molecule expression (P-selectin), increased ischaemia-induced vasoconstriction, during vessel occlusion, and an impaired flow-mediated dilation response of the iliac artery in vivo.

**Conclusions:** In summary, XOR derived NO is critical for maintaining vascular homeostasis under physiological conditions and is key in mediating the benefits of dietary nitrate regimes in cardiovascular pathology

## Introduction

Xanthine oxidoreductase (XOR) is a molybdoflavin enzyme expressed at low level widely but with high expression in the liver^1^. Whilst XOR has been localised to the cytoplasm of various cells of the cardiovascular system^2,3^ it has also been localised to the extracellular surface. Evidence suggests that this surface expression is derived predominantly from the liver, where hepatocyte-derived XOR is secreted into the circulation from where it then readily binds to the surface of endothelial cells^4^, and probably red blood cells (RBCs)^5^, via its interaction with glycosaminoglycans (GAGs) on proteoglycans.

XOR catalyses the oxidation of hypoxanthine to xanthine and xanthine to uric acid (UA). These reactions generate a flow of electrons from the molybdenum site to the flavin adenine dinucleotide (FAD) site, which in the presence of oxygen results in the production of reactive oxygen species (ROS), namely superoxide (O_2_^-^) and hydrogen peroxide (H_2_O_2_). Clinical studies have demonstrated an elevation of XOR activity and expression in patients with hypertension and coronary artery disease^6–8^. In addition, numerous pre-clinical *in vivo* studies have suggested that xanthine oxidase derived ROS are, in part, responsible for the leukocyte recruitment observed in ischaemia reperfusion injury^9^. These data suggest that elevated XOR activity and/or expression is pathogenic, although whether this is due to elevated ROS production or elevated UA generation, which has pro-oxidant activity in disease pathologies^10,11^, remains unclear.

A hallmark of cardiovascular disease is endothelial dysfunction which is synonymous with a lack of bioavailable nitric oxide (NO), in part, due to scavenging by ROS^12,13^. Whilst XOR is well recognised as a ROS generator, evidence indicates that a third function of the enzyme is its capcity to reduce nitrite to NO^14–17^. This activity has been implicated as a key second step in non-canonical (L-arginine-NO synthase independent) NO generation^18,19^. Briefly, inorganic nitrate, found in green leafy vegetables, enters the circulation, concentrating within the salivary glands. Nitrate-rich saliva is excreted into the oral cavity where it is chemically reduced to nitrite by commensal bacteria expressing nitrate reductases^20,21^. This nitrite-rich saliva once swallowed enters the circulation^22^ where it is reduced to NO. A number of distinct nitrite reductases have been identified, however there is growing support for the importance of XOR as a nitrite reductase^23^. Evidence shows that generation of functional NO via the non-canonical pathway occurs in both health and disease however, whilst XOR has been strongly implicated as an effective NO generator in disease, what role it might play as a nitrite reductase in health is largely unknown. Thus, in this study we investigated the role of XOR-dependent nitrite reductase activity in cardiovascular homeostasis utilising *Xdh* transgenic mice. Additionally, to overcome the lethality of homozygous *Xdh* transgenic mice, the limitations of pharmacological XOR inhibitors and to investigate a proposed primary source of systemic XOR, we generated a unique hepatocyte (Alb-Cre) specific XOR-deletion mouse.

## Methods

For expanded methods see online supplement.

### Animal studies

All experiments were conducted according to the Animals (Scientific Procedures) Act 1986, United Kingdom, and approved by the UK Home Office.

### Generation of *Xdh* global and *Xdh*–hepatocyte specific (HXOR KO) knockout mice

*Xdh* knockout mice (*Xdh ^-/-^*) originally generated by Finkel and colleagues^24^, were bred in-house. Embryos were purchased and rederived at MRC Harwell in C57BL/6J mice and *Xdh^+/-^* breeding trios setup. A novel C57BL/6J *Xdh* floxed mouse was generated via CRISPR/Cas9 genome editing (MRC Harwell), and then crossed with C57BL/6J Albumin (Alb) Cre mice (purchased from The Jackson Laboratory) to generate hepatocyte specific XOR KO (HXOR) knockout mice in-house (Figure 1 A-B). Both male and female littermate *Xdh*^+/+^, *Xdh*^+/-,^ *Xdh-*^/-^ and *Xdh^fl/fl^* control / HXOR KO mice were used in all studies.

**Figure 1.**
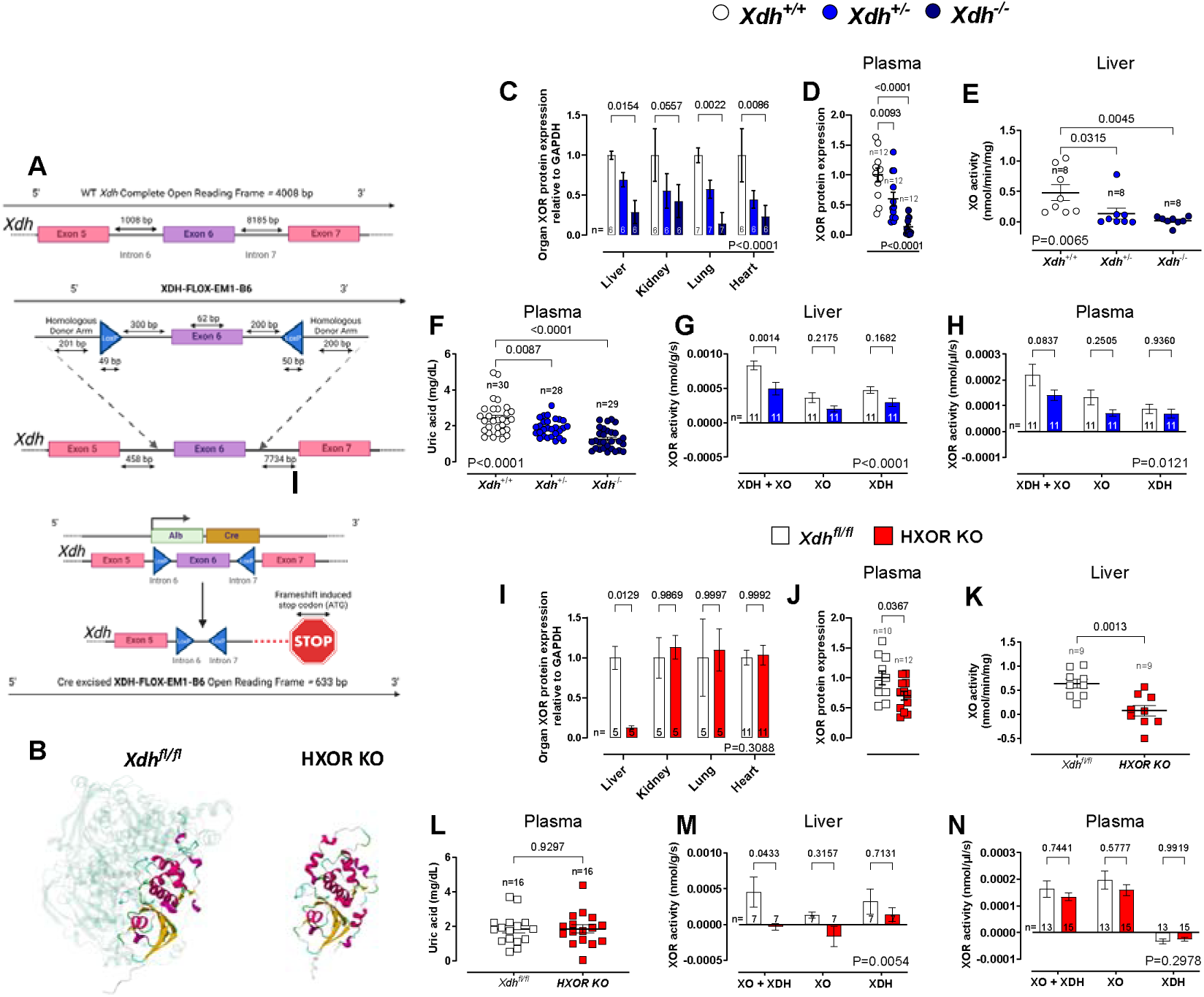
Confirmation of genetic deletion of *Xdh*. **(A)** Schematic of WT *Xdh* gene and insertion of XDH-FLOX-EM1-B6 via CRISPR/Cas9. The expression of the loxP regions in mice containing XDH-FLOX-EM1-B6 (*Xdh^fl/fl^*) results in Cre mediated excision of exon 6 in mice expression Cre recombinase downstream of the albumin promoter (HXOR KO). **(B)** Complete *Xdh* open reading frame in *Xdh^fl/fl^* and HXOR KO mice with 1-633 bp highlighted and integrated into remaining *Xdh* open reading frame protein and formation of an XOR monomer. Western blot quantification of XOR protein expression across **(C)** various tissues and **(D)** plasma in the 4-week-old global *Xdh* deleted mice and their wild type littermate controls. **(E)** XO activity as assessed by amplex red H_2_O_2_ production and **(F)** plasma uric acid concentration and total XOR activity (XDH and XO isoform activity) in the **(G)** liver and **(H)** plasma of samples collected from 4-week-old *Xdh^+/+^*, *Xdh^+/-^* and *Xdh^-/-^* mice where available. Western blot quantification of XOR protein expression across **(I)** various tissues, in 8-week-old and **(J)** plasma in the 20-week-old HXOR KO mice versus the *Xdh^fl/fl^* wild type littermate controls. **(K)** liver homogenate XO activity assessed by amplex red H_2_O_2_ production and **(L)** plasma uric acid concentration and total XOR activity (XDH and XO isoform activity) in the **(M)** liver and **(N)** plasma of 20-week-old *Xdh^fl/fl^* and HXOR KO littermate mice. Data are shown as mean ± SEM of n mice (shown on individual graphs). Statistical significance for genotype was determined using two-way ANOVA (shown in bottom right **(C**, **G-I**, **M-N)** followed by Dunnett’s *post hoc* test **(C and I)** or Sidak’s multiple *post hoc* test **(G-H and M-N),** by One-way ANOVA **(D-F)** followed by Dunnett’s multiple *post hoc* test highlighting significant difference from *Xdh*^+/+^ or by unpaired Student’s t-test **(J-L).**

### Dietary nitrate protocol

Where required, mice (6 weeks old) were randomly assigned to receive drinking water supplemented with potassium nitrate (KNO_3_, 15 mmol/L) or potassium chloride (KCl, 15 mmol/L, control) for 2 weeks^25^. For intravital microscopy studies, mice (3 weeks old) were assigned to the above protocol but for 1 week only.

### Tail cuff blood pressure measurement

Tail cuff blood pressure (BP) data was collected using the non-invasive CODA mouse BP system (Kent Scientific Corporation, USA) and the acquisition conducted blind to genotype.

### Echocardiography

Transthoracic echocardiography was performed using a Vevo 3100 imaging system with a MX550D, 40MHz transducer (FujiFilm VisualSonics Inc., Netherlands). Image analysis was conducted offline and blind to genotype^26^.

### Flow mediated dilation (FMD)

FMD of the arteria iliaca externa was visualized using a Vevo 3100 imaging system with a MX550D, 40MHz transducer (FujiFilm VisualSonics Inc., Netherlands). An O-cuff (Kent Scientific Corporation, USA) was placed above the left hind leg knee and inflated to 300 mmHg at T=0 minutes to occlude blood flow through the left arteria iliaca externa. At T=5 minutes the cuff was released and arteria iliaca externa was visualized until T=10 minutes^27^. Image analysis was conducted offline and blind to genotype.

### Intravital microscopy to determine leukocyte recruitment

4-week-old mice were anaesthetised, the mesentery exposed and superfused with bicarbonate-buffered solution (BBS) and leukocyte rolling and adhesion counted as described previously^25^.

### Flow cytometry

Blood was collected and flow cytometry conducted to determine leukocyte subtypes, activation markers and XOR expression. Additionally, bone marrow was isolated and CXCR2 expression determined.

### Measurement of nitrate and nitrite levels

Plasma and tissue nitrite (NO_2_^-^) and nitrate (NO ^-^) levels (collectively termed NO) were measured using ozone-based chemiluminescence as described previously^25^.

### Liver homogenate nitrite reductase activity

The nitrite reductase activity of liver supernatants was determined using gas-phase chemiluminescence^28^ at pH 7.4 (representing physiological conditions), pH 6.8 (blood acidosis) and pH 5.5 (severe acidosis - conditions that also favour nitrite reductase pathways).

### Immunohistochemistry

Liver and heart samples were collected from paraformaldehyde infused mice, sections prepared and stained with H&E, anti-CD62P, anti-CD45 and picrosirius red. Wheat germ agglutinin conjugated to Alexaflour 647 fluorescent staining was conducted in order to determine cardiac myocyte area^26^.

### Western blotting

Equal amounts of protein homogenates of liver were subjected to Western blotting to determine XOR, phosphorylated (p)-eNOS and total eNOS expression levels.

### Quantitative reverse transcriptase-polymerase chain reaction (qRT-PCR)

Liver and heart samples were subjected to qPCR and analysed using the comparative threshold cycle (Ct) method (2−ΔΔCt). See Table S1 for a full list of primers.

### RNA sequencing analysis

RNA was extracted from cardiac homogenates of global *Xdh^+/+^* and *Xdh^+/-^*mice and subjected to RNA sequence analysis.

### Liver XO activity

XO activity of liver homogenates was determined using a commercially available kit, according to manufacturer guidelines (xanthine oxidase assay kit, Abcam, UK).

### Pterin-based fluorometric assay of XDH/XO activity

To determine the relative proportions of XDH versus XO activity in tissues and plasma we used a modified pterin-based fluorometric assay protocol as previously described^29^.

### Plasma and urine biochemical analysis

Whole blood was obtained via cardiac puncture into a syringe containing 3.8% sodium citrate centrifuged at 14,000 rcf for 5 min at 4 °C and plasma collected. Plasma and spot urine samples were collected at the time of sacrifice, snap frozen and stored at −80°C. Plasma and urine samples were analysed by MRC Harwell Pathology using the AU680 Analyser (Beckman Coulter).

### Sample size estimation

Preliminary exploratory studies were conducted to assess global KO anion levels and/or BP in age- and sex-matched littermate mice. For these experiments formal sample size estimates were not conducted. For all echocardiography studies, power calculations were based on the sample size required to detect a significant difference in left ventricular (LV) posterior wall thickness. In previous studies, we demonstrated that in mice treated with 2 mg/kg/day Angiotensin II, LV wall thickness in KCl placebo control was 1.3 ± 0.4 mm (mean ± SD), compared to 0.81 ± 0.2 mm in mice treated with KNO ^26^ Using these values, with α = 0.05 and a power of 95%, the required sample size was calculated to be n=12 animals per group. To account for potential technical failures or phenotype-related loss of animals, the sample size was increased to n=15 per group for both mouse models. Tail-cuff BP measurements were also performed in *Xdh^fl/fl^* and HXOR KO mice that underwent echocardiography (n = 15). For qPCR, initial assessments of liver TNFα mRNA expression in 4-week-old *Xdh^-/-^* mice yielded a relative expression of 1.0 ± 0.66 (mean ± SD), while in *Xdh^+/-^* mice, relative expression was 2.0 ± 0.77 (effect size 1.38). Using these values and a power of 95%, a sample size of n=15 mice per group was determined to be sufficient for statistical significance. However, for all qPCR and biochemical analyses, all available samples were utilised, on occasion exceeding n = 15. All groups compared were matched for sex, age, and littermate status.

### Statistical and data analysis

Data are expressed as mean ± SEM of n mice. Statistical significance was determined using unpaired t-test, one-way ANOVA followed by Dunnett’s post hoc t-test or two-way ANOVA with Sidak’s post-test as required. A P value <0.05 was considered significant. Gaussian distribution and homogeneity of variances of datasets were determined by the Shapiro-Wilk test and Bartlett tests, respectively. For all experiments n values were equal by design however where unequal n values within experiments are shown this relates to technical failure or from identification of outliers using the ROUT^30^ exclusion test. Where any exclusions have occurred, this has been stated explicitly in the figure legends. All analysis was conducted using GraphPad Prism 10.1.2.

## Results

### Biochemical and molecular characterisation of global and liver specific *Xdh* transgenic mice

As anticipated, typical numbers of offspring and growth rates were observed in the global *Xdh^+/+^ and Xdh^+/-^* mice, while the *Xdh*^-/-^ mice had stunted growth with lethality by ∼4 weeks of age (Figure S1A and S1B)^24^. Consequently, samples from *Xdh*^-/-^ mice were unavailable for non-juvenile (> 4-weeks-old) *in vivo* studies. Whilst weight was similar at 4 and 12 weeks of age, at 20 weeks *Xdh^+/-^* mice were modestly heavier than their *Xdh^+/+^*littermates (Figure S1A). Although when expressed as a change from baseline no statistical differences between *Xdh^+/-^* and *Xdh^+/+^* were evident. In contrast to the *Xdh*^-/-^ mice, there were no survival issues observed in the HXOR KO mice compared to control *Xdh^fl/fl^* littermates (Figure S1C).

Global *Xdh* excision in *Xdh*^-/-^ mice was confirmed by assessment of mRNA (Figure S1D) and protein expression (Figure 1C, 1D and Figure S1F, G) in several tissues and plasma. In contrast, in HXOR KO mice, *Xdh* mRNA (Figure S1E) and XOR protein expression (Figure 1I and Figure S1H) were profoundly reduced in the liver only, whilst expression in other organs remained unaltered. HXOR KO mice had a significant but modest reduction of plasma XOR expression, confirming that hepatocyte XOR contributes to circulating plasma XOR levels (Figure 1J and Figure S1I).

XO activity was significantly reduced in liver homogenates of *Xdh^+/-^* mice versus *Xdh^+/+^* mice and entirely absent in *Xdh^-/-^*mice, (Figure 1E). Similarly, XO activity was reduced in HXOR KO mice in comparison to the *Xdh^fl/fl^*mice (Figure 1K). Interestingly, whilst an allele dependent reduction in plasma UA levels was evident in the global deletion mice (Figure 1F) no such effect was observed in HXOR KO mice (Figure 1L). Assessment of XDH and XO proportions in liver samples demonstrated an equivalent decrease in activity for both isoforms in the global deletion mice (Figure 1G), with similar reductions in plasma XDH and XO levels in *Xdh^+/-^*mice (Figure 1H). In contrast, although a complete ablation of XDH and XO activity was evident in liver samples of HXOR KO mice (Figure 1M), only a modest reduction in plasma activity was evident, which did not reach statistical significance (Figure 1N).

The reductions in XOR expression were not associated with any overt changes in general circulating biochemical parameters measured in 4-week-old global KO mice and 20-week old HXOR KO mice, except for urea levels which were reduced in the global *Xdh^+/-^* mice, but not evident in the liver-selective deleted mice (Table S2) aligning with previous observations^24, 31^. Importantly, neither *Xdh^+/-^* or HXOR KO mice expressed differences in markers of liver damage, including ALP, AST and ALT, compared to their wild type controls (Table S2).

### *Xdh* deletion alters nitrite reductase activity and influences nitrite and nitrate dynamics

To explore whether *Xdh* deletion impacts circulating nitrite and nitrate levels, we measured both anions in tissues and blood of mice of ∼4 weeks of age to enable comparisons between *Xdh^+/+^*, *Xdh^+/-^* and *Xdh^-/-^* littermate mice. We observed an allele-dependent elevation in liver and plasma nitrite and nitrate levels in globally deficient mice (Figure 2A-D). These changes were associated with an allele-dependent decrease in nitrite reductase activity at pH 7.4 and 6.8 in liver homogenates and plasma (Figure 2I–K). Interestingly, in comparison to *Xdh^fl/fl^* mice, HXOR KO mice expressed an increase in liver nitrate levels at 20 weeks (Figure 2F) corresponding with a decrease in plasma nitrite levels (Figure 2G). The age-dependent changes in anion levels, in the HXOR KO mice, were also associated with reduced nitrite-reductase activity in both liver and plasma in 20-week-old mice (Figure 2M-O). To determine whether the reductions in nitrite reductase activity led to deficiencies in vascular NO, we assessed platelet cGMP levels as a marker of NO activity. Indeed, in both the *Xdh^-/-^* and HXOR KO mice platelet cGMP was reduced in comparison to control littermate mice, *Xdh^+/+^* and *Xdh^fl/fl^* mice respectively, although this did not reach statistical significance in HXOR KO mice (Figure 2L and P).

**Figure 2.**
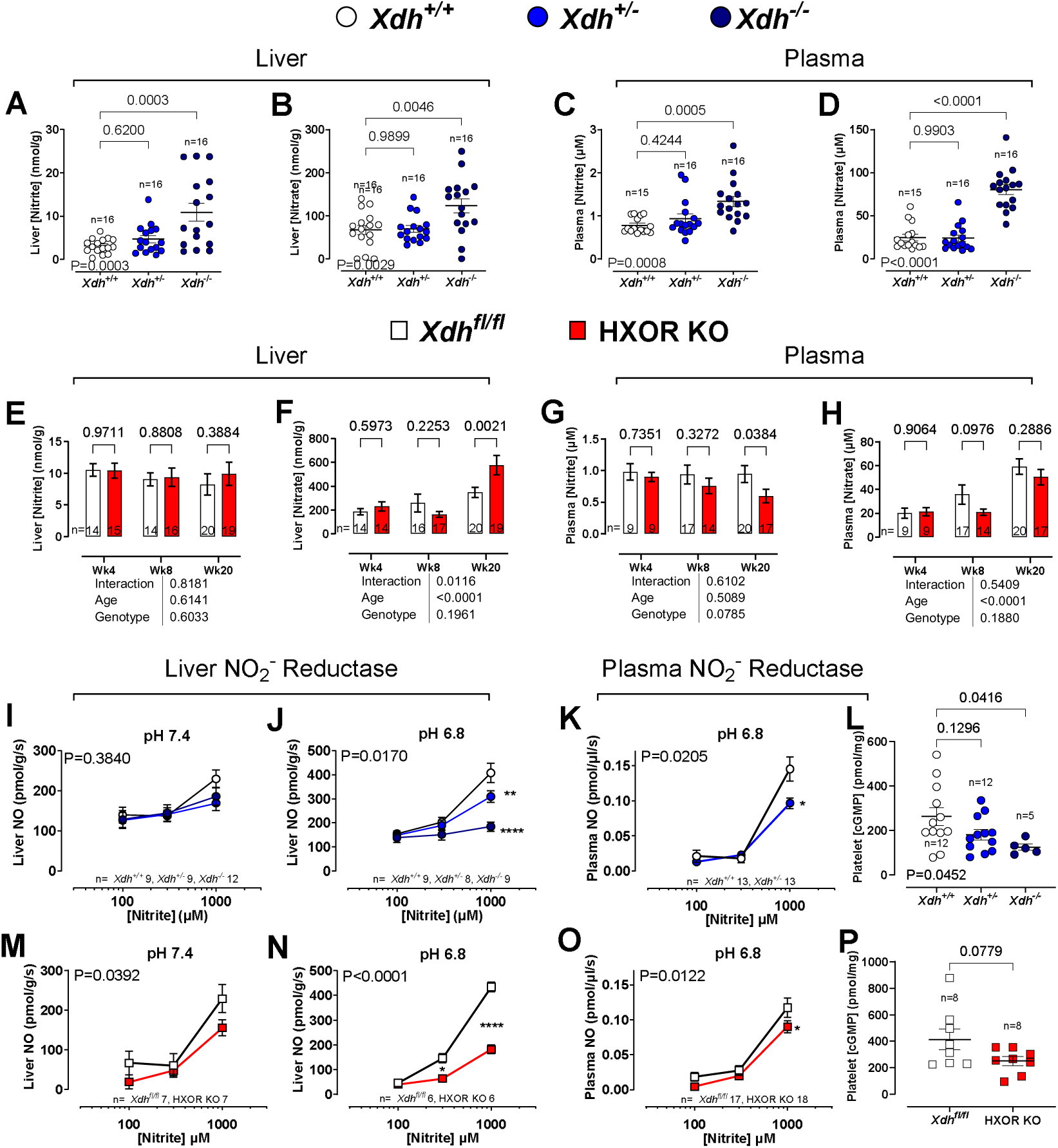
XOR functions as a nitrite reductase. Levels of liver **(A)** nitrite and **(B)** nitrate, and plasma **(C)** nitrite and **(D)** nitrate in 4-week-old global *Xdh* deleted mice and their wild type littermate controls. Levels of liver **(E)** nitrite and **(F)** nitrate, and plasma **(G)** nitrite and **(H)** nitrate in HXOR KO mice versus the *Xdh^fl/fl^* wild type littermate controls. Nitrite reductase activity of liver homogenates at **(I)** pH 7.4, and **(J)** pH 6.8 and **(K)** plasma at pH 6.8 and **(L)** cGMP concentration of platelets isolated from blood samples collected from 4-week-old global *Xdh* deleted mice versus their wild type littermate controls. Nitrite reductase activity in liver homogenates, at **(M)** pH 7.4, and **(N)** pH 6.8 and **(O)** plasma at pH 6.8 **(P)** and cGMP concentration of platelets isolated from blood samples, collected from 20-week-old HXOR KO mice versus the *Xdh^fl/fl^* wild type littermate controls. Data are shown as mean ± SEM of n mice (shown on individual graphs). Statistical significance was determined using one-way ANOVA followed by Sidak’s multiple post-tests highlighting significant difference from *Xdh*^+/+^ **(A-D and L),** using mixed effect analysis **(E-H),** two-way ANOVA **(M-O)** followed by Sidak’s multiple *post hoc* tests shown as *P<0.05, **P<0.01, ***P<0.001 vs *Xdh^+/+^* or *Xdh^fl/fl^* at respective concentration, or unpaired Student’s t-test **(L and P)**.

There is some suggestion that in the absence of functional XOR, compensatory changes may occur in eNOS expression and/or activity. To assess whether changes in eNOS might account for the above noted effects we assessed total eNOS mRNA (Figure S2A and S2B) and protein expression (Figure S2D, S2F, S2G and S2I), as well as phospho-eNOS expression (Figure S2E and S2H) as a measure of activity, in liver homogenates. We found no evidence of differences in the *Xdh^-/-^* and HXOR KO mice versus the control littermate *Xdh^+/+^* and *Xdh^fl/fl^* mice respectively.

### Hepatocyte *Xdh* deletion elevates BP and promotes cardiac remodelling

Since there is substantial evidence indicating that the non-canonical pathway for NO delivery can profoundly influence BP we measured BP in both mouse models over 24 weeks. However, since *Xdh^-/-^* die at ∼4 weeks of age, only *Xdh^+/+^* and *Xdh^+/-^* mice were available. At 4 weeks of age *Xdh^+/+^* and *Xdh^+/-^* mice exhibited comparable BP. However, in agreement with previous findings^30^ between 8-24 weeks of age *Xdh^+/-^* mice exhibited elevated systolic BP (SBP), diastolic BP (DBP) and mean arterial pressure (MAP) versus litter and age-matched *Xdh^+/+^* mice (Figure 3A-C). Importantly, this phenotype was replicated in mice with hepatocyte selective deletion of *Xdh,* with HXOR KO mice expressing elevated SBP, DBP and MAP from 10 weeks of age (Figure 3D-F).

**Figure 3.**
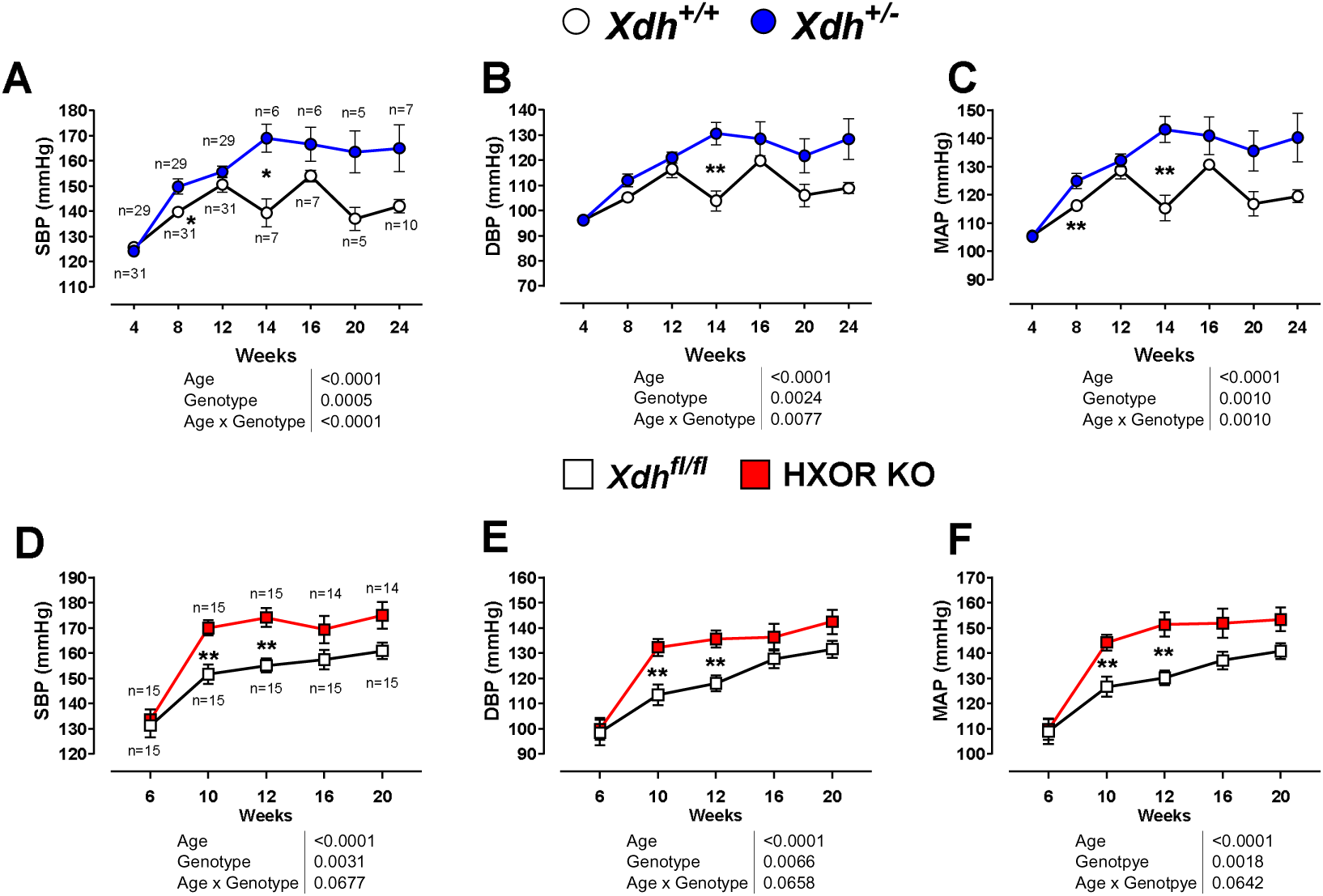
*Xdh^+/-^*and HXOR KO mice exhibit raised blood pressure. **(A)** Systolic, **(B)** diastolic, and **(C)** mean arterial pressure were raised in *Xdh^+/-^* versus *Xdh^+/+^* mice and **(D-F)** HXOR KO versus *Xdh^fl/fl^* mice. Data are shown as mean ± SEM of n mice (shown on graphs **(A)** and **(D)**). Statistical significance was determined using mixed-effect analysis followed by Sidak’s multiple *post hoc* tests shown as *P<0.05, **P<0.01, ***P<0.001.

Since the BP phenotype was evident between 8-10 weeks, we conducted echocardiography from this timepoint onwards. Overall, Figure 4 demonstrates that *Xdh^+/-^* mice develop significant LV hypertrophy (Figure 4A), at 16 weeks of age, following the onset of hypertension. The LV diastolic volume (Figure 4B) and area (Figure 4E) are significantly enlarged however, the diastolic internal diameter of the LV (Figure 4F) is not affected. This LV hypertrophy is accompanied by significant thickening of the LV posterior wall (Figure 4I) in *Xdh^+/-^* mice. No significant changes in LV systolic parameters were observed in *Xdh^+/-^* mice in either the short axis (Figure S3) or parasternal long axis (Figure S4). This compensatory LV diastolic wall thickening in *Xdh^+/-^* mice likely prevents a reduction in ejection fraction (Figure 4M) and stroke volume (Figure 4N), resulting in *Xdh^+/-^* mice having a cardiac output (Figure 4Q) comparable to that of *Xdh^+/+^* littermates.

**Figure 4.**
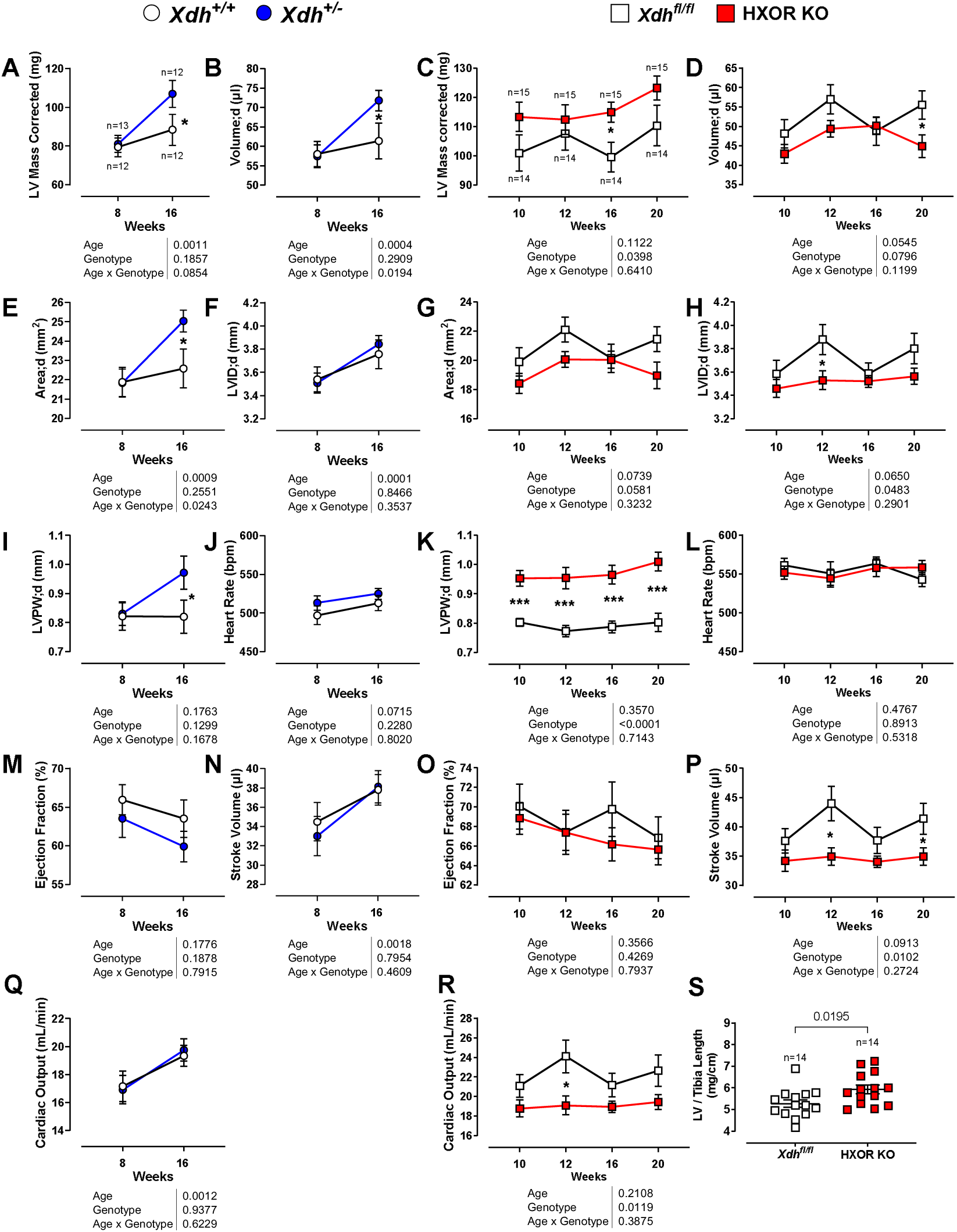
*Xdh^+/-^*and HXOR KO mice exhibit left ventricular remodelling. Echocardiography data of *Xdh^+/-^* versus *Xdh^+/+^* littermates, at 8-16 weeks, and *Xdh^fl/fl^* versus HXOR KO littermates at 10-20-weeks-old. **(A)** Short axis M-mode (SAX) LV corrected mass, and **(B)** parasternal long axis B-mode (PSLAX) acquisition of LV volume in diastole of *Xdh^+/-^* versus *Xdh^+/+^* littermates. **(C)** SAX LV corrected mass and **(D)** PSLAX LV volume in diastole of HXOR KO versus *Xdh^fl/fl^*. **(E)** PSLAX LV area in diastole, **(F)** SAX LV internal diameter in diastole, **(G)** PSLAX LV area diastole, **(H)** SAX LV internal diameter in diastole, **(I)** SAX LV posterior wall thickness in diastole, **(J)** SAX heart rate, **(K)** SAX LV posterior wall thickness in diastole, **(L)** SAX heart rate, **(M)** SAX ejection fraction, **(N)** SAX stroke volume, **(O)** SAX ejection fraction, **(P)** SAX stroke volume, **(Q and R**) SAX cardiac output, and **(S)** dissected LV and tibia length from 20-week-old HXOR KO mice versus *Xdh^fl/fl^*littermates. Data are shown as mean ± SEM of n mice (shown on graphs **(A)** and **(C)**). Statistical significance w*^-^*as determined using mixed-effects or Two-ANOVA analysis followed by Sidak’s multiple *post hoc* tests at an individual timepoint shown as *P<0.05, **P<0.01, ***P<0.001.

Similarly, HXOR KO mice display significant LV hypertrophy coinciding with the onset of hypertension (Figure 4C). However, in contrast to the global KO model, the remodelling in HXOR KO mice appeared to be less compensatory and more maladaptive. HXOR KO mice displayed a trend of reduced LV volume (Figure 4D) and area (Figure 4G) in both diastole and systole (Figure S6), which corresponds to a significant decrease in total LV volume (Figure S6H) and area (Figure S6F). This smaller LV area is attributed to a significant reduction in the diastolic internal diameter (Figure 4H). The reduced internal diameter in HXOR KO mice may be due to significant thickening of both the posterior (Figure 4K and S5K) and anterior walls (Figures S5E and S5F). The pathological LV remodelling and reduced internal diameter in HXOR KO mice did not affect heart rate (Figure 4L) or ejection fraction (Figure 4O) but did significantly attenuate LV stroke volume (Figure 4P), which in HXOR KO mice corresponds with a decreased cardiac output (Figure 4R). To further assess LV mass, excised LVs were collected and weighed from 20-week-old HXOR KO and *Xdh^fl/fl^* littermate mice. Figure 4S demonstrates that, in accordance with the echocardiographic data, the LV mass normalised to the tibia length of HXOR KO mice is significantly larger than the wild type controls (Figure 4S).

Further assessment of diastolic function in HXOR KO mice revealed no single parameter showing a statistically significant change. However, a trending increase in the E/E’ ratio (Figure S8H), LV myocardial performance index (MPI) using isovolumetric parameters (IV) (Figure S8I), and LV MPI using non-filling time (NFT) (Figure S8J) were evident. The impaired cardiac output in HXOR KO mice also corresponds with a trend towards reduced aortic velocity time integral (VTI) (Figure S9A) relative to their littermate controls.

Since *in vivo* functional studies suggested BP-induced changes in cardiac function occurred as early as 8 weeks of age and increased thereafter, we measured immunohistochemical markers of hypertrophy and/or fibrosis at both 8 and 20 weeks of age. At 8-weeks of age *Xdh^+/+^* and *Xdh^+/-^*mice expressed comparable collagen content and cardiomyocyte area (Figure 5B and C), and a similar pattern was also observed in 8-week-old HXOR KO mice vs *Xdh^fl/fl^* (Figure 5D and E). However, in 20-week-old HXOR KO mice when both raised BP and cardiac dysfunction are strongly expressed, elevated collagen deposition was observed within the heart (Figure 5D) associated with a trend towards increased cardiomyocyte area (Figure 5E).

**Figure 5.**
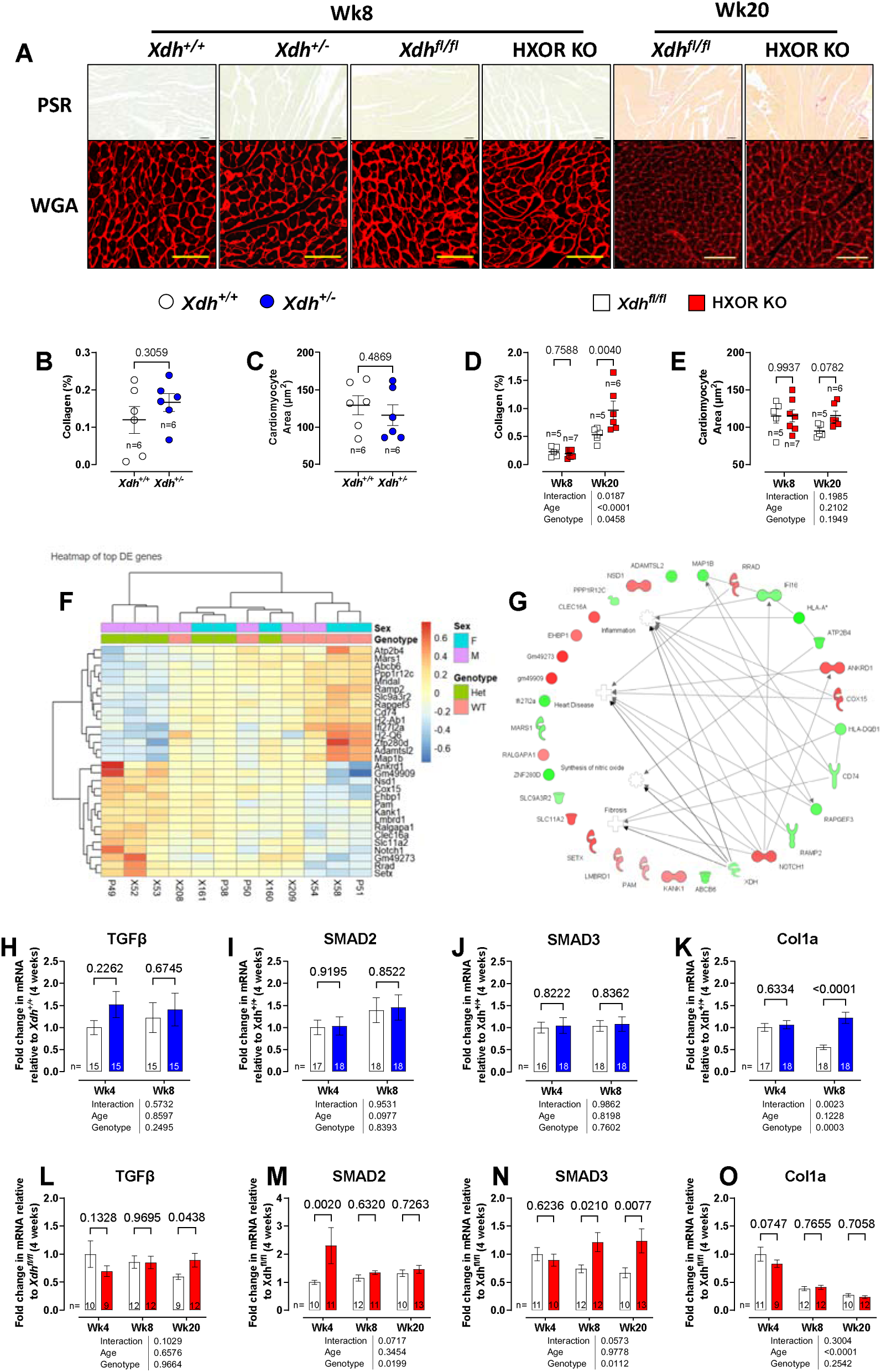
Upregulation of profibrotic pathways in *Xdh^+/-^* and HXOR KO mice. **(A)** Representative images of picrosirius red (PSR) and wheat germ agglutinin staining (WGA) in sections of formalin fixed left ventricle (LV). Quantification of levels of **(B, D)** cardiac collagen deposition and **(C, E)** cardiomyocyte area in both the global *Xdh^+/-^* and HXOR KO mice versus littermate controls. **(F)** Heatmap summary of all differentially expressed (DE) genes (q<0.1) identified in total mRNA extracted from whole hearts of 8-week-old *Xdh^+/+^* and *Xdh^+/-^* mice. **(G)** Network plot demonstrating key disease and biological function annotations across DE genes. Longitudinal mRNA expression profiling in total mRNA extracted from whole hearts of *Xdh^+/-^* and HXOR KO mice versus littermate controls investigating markers of fibrosis: **(H, L)** TGFβ, **(I, M)** SMAD2, **(J, N)** SMAD3, and **(K, O)** Col1a. Data are shown as mean ± SEM of n mice (shown on individual graphs). Statistical significance was determined using mixed effects analysis followed by Sidak’s multiple *post*-*hoc* tests. **(D-E, H-O)** or unpaired Student’s t-test **(B, C).** Uneven n values relate to sample loss due to technical failure or Routs exclusion.

To explore the potential signalling pathways that might drive these effects we conducted bulk RNA sequencing of cardiac homogenates of 4-week-old mice to better explore the initiating signalling pathways triggered by an absence of *Xdh.* Our analyses demonstrate an upregulation of pro-fibrotic pathways in the young *Xdh^+/-^* mice, indicative of the triggering of cardiac remodelling compared to their *Xdh^+/+^* littermate controls (Figure 5F and G). Accordingly, qPCR of key cardiac pro-fibrotic signalling pathways at 4-20 weeks suggested some (albeit modest) activation of the TGFβ−SMAD-Col1a pathway with a trend to increase in TGFβ expression in the 4-week-old *Xdh^+/-^*mice and increased Col1a at 8 weeks (Figure 5H-K) with evidence of upregulated SMAD2-3 in the HXOR KO mice relative to their *Xdh^fl/fl^*littermates (Figure 5L-O). These findings hint at the suggestion that any pro-fibrotic effects in the heart were likely due to an absence of circulating XOR rather than XOR expressed in cardiac cells. Collectively these data add further support to the view that impaired liver XOR-dependent nitrite reductase activity in HXOR KO mice results in a phenotype resembling that of early heart failure with a preserved ejection fraction (HFpEF).

### Reduced hepatic nitrite reductase activity leads to systemic inflammation and endothelial dysfunction

Interestingly, bulk RNA seq analyses of whole heart homogenates also highlighted the upregulation of pro-inflammatory signalling pathways in *Xdh^+/-^* mice (Figure 5G). To explore this further we conducted qPCR of cardiac supernatants in both the *Xdh^+/-^*and HXOR KO mice relative to their littermate controls for a range of proinflammatory mediators (Figure S10 and S11). These results reveal mild upregulation of pro-inflammatory mediators in the hearts of aging HXOR KO mice, particularly TNFα (Figure S10G and S11G).

In cardiovascular disease, systemic inflammation drives endothelial dysfunction that often precedes cardiac dysfunction. Endothelial NO deficiency plays a critical role in this process. Therefore, we assessed endothelial function *in vivo*. As expected, cuff inflation to occlude the external iliac artery followed by cuff release, led to a prominent increase in vessel diameter due to shear stress-induced endothelial activation in control *Xdh^+/+^* mice (Figure 6A). Interestingly, whilst there was minimal change in the vessel diameter during the occlusion period (1-5 min) in *Xdh^+/+^* mice, in *Xdh^+/-^* mice there was a significant decrease in vessel diameter (Figure 6B). After cuff release, *Xdh^+/-^* mice demonstrated significantly reduced vessel dilation compared to *Xdh^+/+^* mice (Figure 6A and C). In HXOR KO mice a similar response profile to both occlusion (vasoconstriction) and shear stress (reduced FMD) relative to their littermate controls was observed (Figure 6D-F). These findings suggest that hepatocyte XOR plays a crucial role in maintaining systemic endothelial function and blood flow.

**Figure 6.**
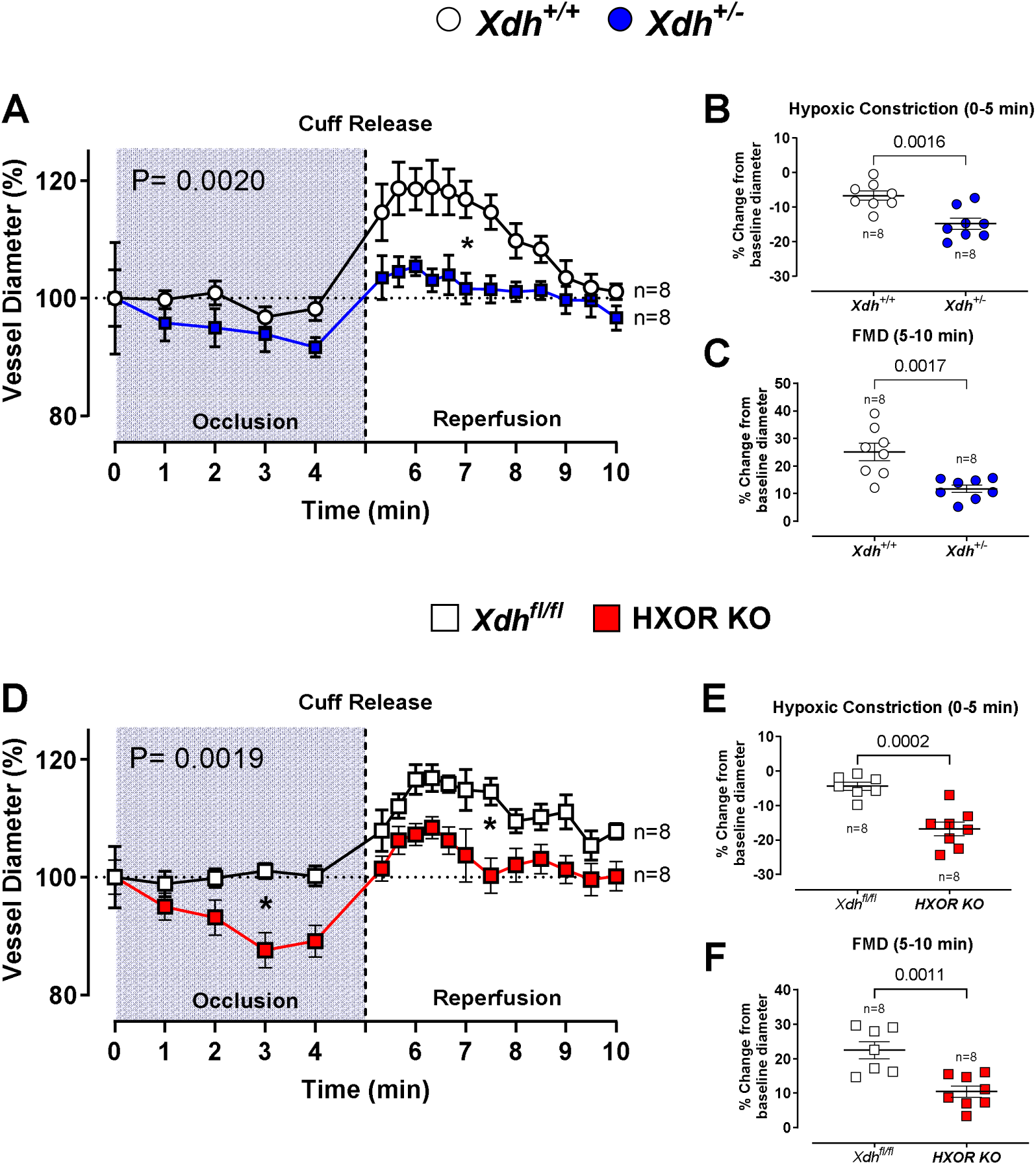
HXOR KO and *Xdh*^+/-^ mice have impaired flow mediated dilatation (FMD). FMD was assessed in both 20-week-old **(A)** *Xdh^+/-^* and (**D)** HXOR KO mice versus littermate controls. An occlusion cuff placed above the left hind leg knee was inflated to 300 mmHg at T= 0 minutes to occlude blood flow through the left arteria iliaca externa. Time frame of occlusion is represented by the blue box. At T= 5 minutes the cuff was released (represented by the vertical dotted line). **(B and E)** show the maximal constriction of the vessel recorded during the occlusion phase expressed as a percentage change from the baseline diameter. The maximal FMD recorded post cuff release **(C and F)**. Data are shown as mean ± SEM of n mice (shown on individual graphs). Statistical significance was determined using two-way ANOVA followed by Sidak’s multiple *post-hoc* tests **(A, D)** with **p***ost hoc* test at an individual timepoint shown as *P<0.05, or using unpaired Student’s t-test **(B, C, E, F)**.

### XOR-deficiency triggers hepatic inflammation driving systemic inflammation and endothelial dysfunction

Since the mild inflammatory effects were observed in both the liver-specific knockout and the globally deleted mice, we hypothesized that the cardiac dysfunction and mild inflammation in the heart might be secondary to a systemic inflammation that originated within the liver of these animals. qPCR analyses of key inflammatory mediators revealed increased mRNA expression of TNFα, particularly at young ages, in the liver of both *Xdh^+/-^* (Figure 7A) and HXOR KO mice relative to their wild type littermates (Figure 7C). We confirmed these changes in TNFα expression in the plasma of 4-week-old *Xdh^-/-^* mice (Figure 7B) and a trend to increase in 20-week-old HXOR KO mice (Figure 7D). No statistically significant alterations were observed in other surveyed inflammatory mediators, except for NLRP-1 and TLR5 in HXOR KO mice (Figures S12, S13, and S14).

**Figure 7.**
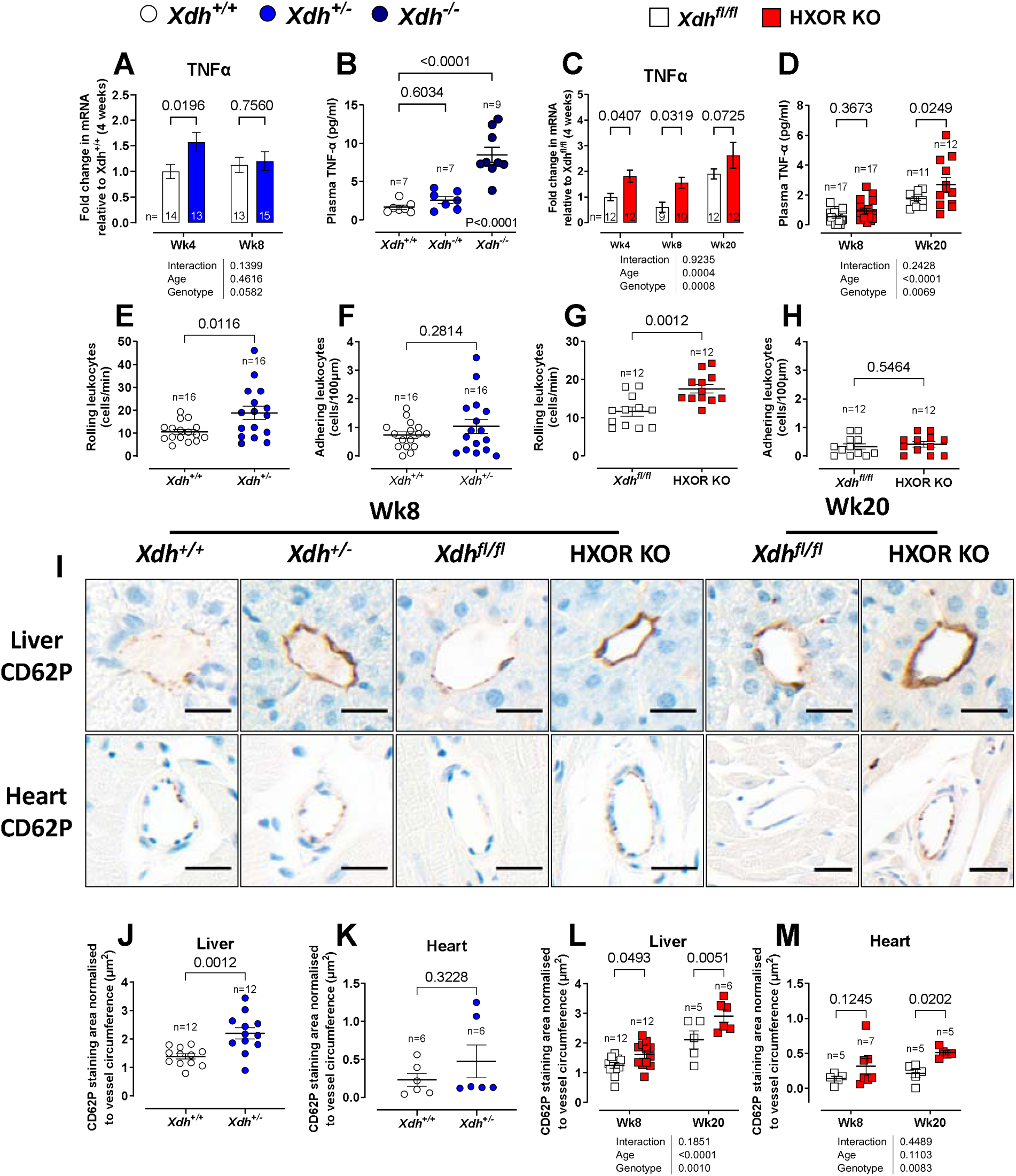
*Xdh^+/-^* and HXOR KO mice exhibit elevated systemic inflammatory state. Longitudinal TNFα mRNA expression in liver homogenate and plasma TNFα levels in 4-week-old of *Xdh^+/-^* mice versus *Xdh^+/+^* littermates (**A-B)** and HXOR KO mice versus *Xdh^fl/fl^* littermates **(C)** and **(D).** Brightfield intravital microscopy was used to determine the number of **(E and G)** rolling and **(F and H)** adhering leukocytes in mesentery vessels of 4-5-week-old *Xdh^+/-^* and HXOR versus wild type littermate mice. Representative images of CD62P staining (**I**) in vessels of liver and heart and their quantification **(J and L)** and **(K and M)** of *Xdh^+/-^* and HXOR versus wild type littermate mice. Data are shown as mean ± SEM of n mice (shown on individual graphs). Statistical significance was determined using two-way ANOVA followed by Sidak’s multiple *post-hoc* tests **(A, C, L, and M)** or using unpaired Student’s t-test **(B, D, F-K).** Uneven n values relate to technical failure or exclusion using ROUTS.

Confirmation of the functional impact of an increased systemic inflammatory state and dysfunctional endothelium was obtained from intra-vital microscopy studies. In 4–5-week-old mice, leukocyte rolling was enhanced in *Xdh^+/-^* compared to *Xdh^+/+^* mice (Figure 7E), while leukocyte adhesion remained comparable between the groups (Figure 7F). Similarly, HXOR KO mice exhibited increased leukocyte rolling (Figure 7G) with no change in adhesion (Figure 7H) relative to the *Xdh^fl/fl^* littermates.

Interestingly, there were no observed differences in the number of CD45+ cells within the liver or heart of 8 or 20-week-old *Xdh^+/-^*or HXOR KO mice (Figure S15 and S16). However, indicative of a nascent pro-inflammatory state and emerging cardiac dysfunction due to deficient NO generation, there was an increase in the expression of the NO-sensitive endothelial adhesion molecule CD62P (P-selectin) in endothelial cells of both the liver and heart of *Xdh^+/-^* and HXOR KO mice (Figure 7J-M). That the enhanced leukocyte rolling is due primarily to changes in endothelial activity rather than direct changes in leukocyte activity is supported by the lack of difference in circulating leukocyte number (Table S4) or activation state (Table S5) between *Xdh^+/-^* and *Xdh^+/+^*mice. These findings suggest the pivotal role of hepatocyte-derived XOR in triggering hepatic inflammation that initiates a systemic pro-inflammatory response leading to generalised endothelial dysfunction, elevated BP, and ultimately cardiac remodelling.

### *Xdh* deletion prevents dietary nitrate-induced protection against ischaemia-reperfusion injury

We investigated the impact of XOR deletion on ischaemia/reperfusion (I/R) injury-induced leukocyte activation and recruitment, given the previous implication of XOR in this response. As expected *Xdh^+/+^* mice exhibited elevated levels of leukocyte rolling (Figure 8A) and adhesion (Figure 8B) in response to I/R injury compared to basal conditions. However, these effects were absent in *Xdh^+/-^* mice, with only a small elevation in leukocyte adhesion observed under I/R injury conditions (Figure 8A and B). These findings suggest that while XOR is protective under physiological conditions, it contributes to leukocyte recruitment during I/R injury. In addition, in *Xdh^+/+^* mice, treatment with inorganic nitrate intervention resulted in a trend to reduction in I/R-induced leukocyte rolling (Figure 8C) with no change in adhesion (Figure 8D). In contrast this intervention had no effect in *Xdh^+/-^*mice. Analysis of circulating leukocyte numbers in mice supplemented with inorganic nitrate showed a statistically significant reduction in neutrophil and inflammatory monocyte counts in *Xdh^+/+^* mice, which was attenuated in *Xdh^+/-^* mice, with no changes observed in other leukocyte subtypes (Table S6). To assess if this was due to neutrophil mobilisation, as a surrogate for this we assessed CXCR2 expression on circulating and bone marrow neutrophils and inflammatory monocytes, finding no significant changes with inorganic nitrate treatment (Table S7). Moreover, the expression of CD162, CD62L and CD11b on various leukocyte populations remained unaltered with inorganic nitrate intervention (Table S8). Collectively, these data suggest that XOR plays a functional role in mediating leukocyte recruitment under physiological conditions and may also help attenuate I/R-induced leukocyte recruitment in the presence of increased circulating nitrite.

**Figure 8.**
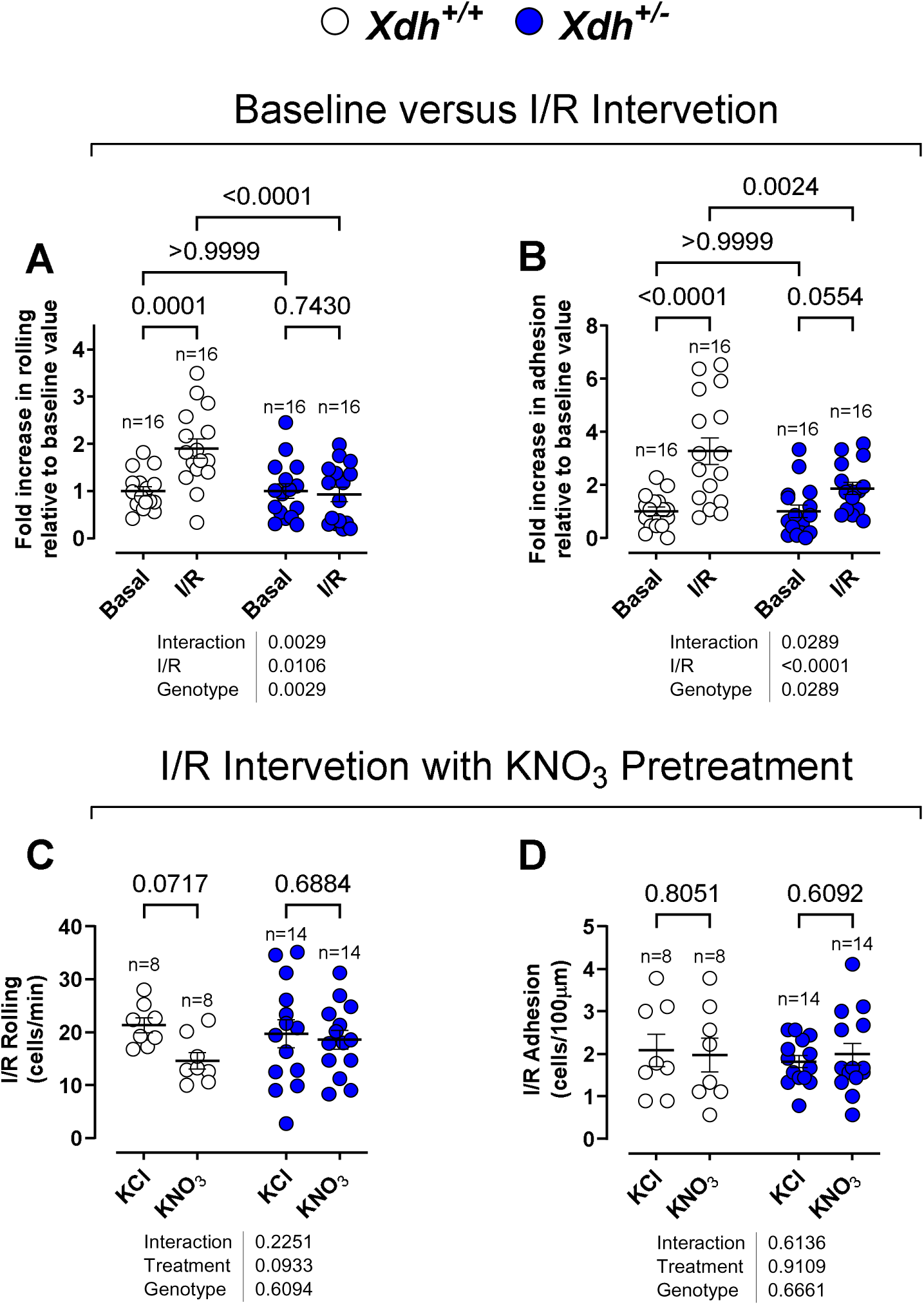
XOR enhances leukocyte recruitment under I/R injury, which is reduced with inorganic nitrate. Brightfield intravital microscopy was used to visualise leukocyte **(A)** rolling and **(B)** in mesentery vessels following 30 minutes of localised ischemia and 45 minutes of reperfusion in 4-5-week-old *Xdh^+/-^*versus wild type littermate mice. At 3-4 weeks of age drinking water was supplemented with either 15 mM KNO_3_ (nitrate intervention) or 15 mM KCl (vehicle) for 1 week prior to I/R injury. At the end of the 1-week KNO_3_ / KCl pretreatment I/R injury was induced, as previously stated and leukocyte **(C)** rolling and **(D)** adhesion was assessed. Data are shown as mean ± SEM of n mice (shown on individual graphs). Statistical significance was determined using Two-way ANOVA followed by Sidak’s multiple *post-hoc* tests.

Given the observed modulation in leukocyte recruitment, both at baseline and following nitrate intervention, we also investigated whether circulating leukocytes express XOR. We did not detect extracellular XOR expression on any leukocyte subtype in *Xdh^+/+^* or *Xdh^+/-^* mice (Table S9). In contrast, intracellular XOR expression was detected in all leukocyte subtypes assessed, albeit with varying degrees of expression (Table S9). In *Xdh^+/+^* mice, neutrophils exhibited the greatest level of XOR expression, followed by monocytes, lymphocytes and B cells. In *Xdh^+/-^* mice, a similar expression profile was observed, albeit the level of XOR expression in neutrophils was significantly reduced. These data suggest that leukocytes, in particular neutrophils, have the capacity to synthesise XOR, implying their ability to reduce nitrite to NO.

## Conclusion

Herein, we demonstrate a critical role for hepatic XOR mediated bioconversion of nitrite to NO in maintaining vascular and cardiac health. This XOR-derived NO suppresses leukocyte activation, maintains healthy BP and cardiac function. We also demonstrate that XOR may be harnessed to reduce leukocyte activation in conditions such as I/R injury by altering the balance of XOR substrate availability in favour of NO production, through elevation of circulating nitrite levels. Collectively these findings implicate liver-derived XOR in sustaining cardiovascular physiology and, moreover, challenge the view that the function of XOR in the cardiovascular system is solely one of driving disease pathology^31–33^. Moreover, these findings highlight the potential in targeting hepatic XOR through dietary nitrate delivery to prevent vascular damage by attenuating the systemic inflammation that drives vascular dysfunction and cardiovascular disease.

Using the global *Xdh* deletion mouse model^24^ we confirmed previous observations indicating that reduced expression of XOR leads to reductions in the products of its activity i.e. UA and O_2_^-^/ H_2_O_2_. However, in addition we show, for the first time, that this expected reduction in redox activity is accompanied by a clear allele-dependent reduction in nitrite reductase activity in tissues and blood, also reflected by alterations in nitrite and nitrate levels. Using the pterin fluorometric assay, we observed a greater proportion of XOR in the XDH form, vs XO, in the liver, whilst in plasma the proportions were comparable. This difference has been attributed previously to sulfhydryl oxidation once released into the circulation^36, 34^. Since we identified nitrite reductase activity in both the liver and plasma, this observation suggests that both isoforms likely contribute to physiological levels of NO, despite some suggestions that this is not the case^35^.

To further explore the role of XOR-derived NO in cardiovascular function we created a floxed mouse, using a CRISPR-Cas9 approach, to overcome the difficulties associated with global deletion^36^. We focussed specifically upon deletion of hepatocyte *Xdh* since the liver is a primary source of circulating XOR implicated in driving the pathogenesis of cardiovascular disease^31,34,37^. This mouse is different from the recently published mouse model created by Kelley and co-workers using a Neo cassette approach^38^. Basic phenotyping of XOR expression and activity confirmed selective hepatic deletion and showed considerable similarity with the previously published mouse model. However, some important differences between the models were identified. Whilst in the previous study an approximate 50-60% reduction in plasma XOR activity was evident^38^, in this CRISPR-Cas9-induced deletion mouse this was not the case; a finding further supported by our observations that there was no significant change in plasma UA level. This discrepancy between the models may relate to the method used to quantify XOR activity, perhaps off-target deletion, but also possibly due to the method for blood collection. We collected blood using citrate as an anticoagulant to prevent heparin-mediated scavenging of GAG bound XOR found on RBCs or endothelial cells^23,39^. Indeed, our analysis shows that XOR expression was unaffected in other tissues collected from the HXOR KOs versus the littermate controls indicating hepatocyte selectivity. But we also suggest that, in addition to the liver, circulating XOR levels may arise from release by other high XOR-expressing organs, such as for instance the lung or adipose tissue, and thus compensating, at least in part, for the absence of liver-derived circulating XOR^40^, at least in this knockout setting.

In the HXOR KO mice, we identified alterations in NO_x_ levels with a rise particularly in nitrate levels in the liver. We suggest that the elevation in liver and plasma nitrite and nitrate, in both models, is due to reduced XOR-dependent conversion of nitrite to NO. We speculate that this leads to an accumulation of plasma nitrite due to less utilisation/metabolism. Nitrite has a short half-life, primarily due to its oxidation to nitrate, and the rise in nitrate levels likely reflects re-direction of nitrite down this route of metabolism^41,42^. We also speculate that the attenuation of plasma nitrite in the HXOR KO animals is associated with decreased systemic NO bioavailability and oxidation into nitrite since we demonstrate clear reductions in the levels of platelet cGMP. It is noteworthy that in the *Xdh^-/-^* animals renal function declines rapidly as they age due to the build-up of purine crystals in the kidney^24,36^, and thus it is possible that disordered renal clearance contributes in some way to the alterations of NOx levels. However, since there is no evidence for renal dysfunction/failure in either the global heterozygotes or those with selective hepatocyte deletion, it is unlikely that failure of renal clearance underlies the changes seen. These findings support the suggestion that hepatocyte XOR is not the exclusive source of circulating physiological XOR, and that XOR released from other organs into the circulation express nitrite reductase activity. These studies also support the many *in vitro* observations implicating XOR, using pharmacological inhibitors i.e. allopurinol and febuxostat, demonstrating nitrite reductase activity in tissues collected primarily from ‘cardiovascular disease’ scenarios in both experimental animals and patients (for review^18,19,43^). Herein, we now demonstrate that, in contrast to some recent suggestions^44^, in addition to acting as a nitrite reductase in disease, that the nitrite reductase activity of XOR activity also plays an important role in physiology.

Due to the critical role of vascular NO in BP control, we assessed whether XOR deletion influences BP. Both models displayed increasing BP with age versus their respective littermate controls, despite normal BP at 4 weeks. This is in accord with a previous study showing modest SBP elevation at 2 months in *Xdh^+/-^* mice, increasing at 4 and 18 months^30^. Importantly, in our hands this raised BP was associated with endothelial dysfunction; as reflected by impaired FMD response. Since this effect was in the absence of circulating UA changes but was associated with decreased systemic XOR nitrite reductase activity and NO bioavailability, and since reduced endothelium-derived NO bioavailability^45,46,47^ is thought to underlie this effect, we suggest that the hypertension observed in the genetic models is attributed to decreased circulating liver-derived XOR-dependent NO generation at the level of the endothelium.

The mechanism by which XOR influences vascular tone and thus BP has long been debated but remains uncertain. Hyperuricemia has been associated with hypertension^47^, however, in contrast, a recent study has suggested that plasma XOR activity, not serum UA, is positively associated with BP^49^. Additionally, whilst lower levels of oxidative stress were associated with BP, this association was absent at higher oxidative stress levels^48^. There are also meta-analyses suggesting that XOR inhibition lowers BP^49^. But it should be noted that there are also studies showing no effect on BP^50,51^. Collectively, these observations highlight the current confusion regarding the role of XOR on BP. We suggest that this complexity arises from a lack of consideration of the full spectrum of XOR activity. Our data demonstrates that a single allele deletion of *Xdh* results in an elevation of BP, contrasting with suggestions that pharmacological XOR inhibition results in a small but significant lowering of BP in certain patient cohorts^52,53^. A potential explanation for these conflicting data is that inhibition of XOR will result in a lowering of UA and ROS production but also a loss of XOR mediated nitrite reduction. It has been suggested that in a scenario where conventional endothelial NO generation is prevented, i.e. the eNOS KO mouse, that there is a compensatory rise in XOR activity, to restore NO bioavailability^54^. This could lead to speculation that any apparent changes in levels of NO or its metabolites may be related to alterations in other sources of NO^55^. This possibility, however, is unlikely in the current setting since neither eNOS nor phospho-eNOS expression were altered in either genetic model.

Interestingly, our findings also suggest that the normal response of the endothelium to blood flow is, at least in part, dependent upon vascular nitrite reductase activity of XOR. Indeed, occlusion of the iliac artery resulted in pronounced vasoconstriction in XOR-deletion mice, an effect not observed in wild-type controls of either genotype. This indicates that vascular XOR activity is regulated by shear stress and suggests that, under conditions of reduced oxygen tension (which enhances XOR-derived nitrite reductase activity^14,28^ but impairs eNOS-derived NO generation due to its oxygen dependence^54^), XOR compensates by delivering NO. Importantly, we also show that this XOR dependency relates to liver-derived, and likely, endothelial bound, XOR using the HXOR KO mice. Recently, Cortese-Krott and co-workers, demonstrated that the FMD response of the iliac artery is completely absent in endothelial cell specific eNOS-ablated mice. However, these mice did not show any evidence of enhanced vasoconstriction with occlusion^27^. We speculate that upregulation of XOR-dependent nitrite reductase activity in the low oxygen environment underlies this effect. Indeed, there is evidence that eNOS and XOR operate in tandem to ensure vascular NO delivery, and that in the absence of eNOS there is a compensatory increase in XOR-dependent NO generation to maintain vascular homeostasis^54^.

Prolonged elevated BP impacts cardiac function, and indeed, in the global deletion mice, raised BP precedes the cardiac phenotype. However, in HXOR KO mice, we observed a reduction in cardiac output alongside increased left ventricular posterior wall (LVPW) thickening and decreased left ventricular internal diameter (LVID). The disparity in cardiac phenotypes—compensatory LV hypertrophy in the global heterozygous KO model, versus maladaptive remodelling in the homozygous hepatocyte KO model, likely highlights the distinct role of hepatocyte-derived XOR in cardiovascular homeostasis. The HXOR KO mice developed maladaptive cardiac -remodelling as early as 6 weeks of age, before the apparent onset of hypertension, indicating a potential BP-independent component. Indeed, our previous studies^26^ have identified both BP-dependent and independent pathways underlying beneficial effects of activating the non-canonical pathway, through dietary nitrate delivery, in the setting of a severe cardiac dysfunction phenotype. Both pathways are dependent upon NO activity, but the latter is mediated by NO-driven suppression of the TGFβ-SMAD-Col1a pathway of fibrosis. Indeed, this latter possibility likely explains, at least a component of the effects, since upregulated mRNA expression for both TGFβ and Col1a were evident in the hearts of both *Xdh*-deficient models. Importantly, the cardiac changes, in both models, are consistent with a heart failure phenotype with diastolic dysfunction. These observations are in contradiction to a recent study using the Neo cassette approach to hepatic XOR deletion, where no differences in LV structure or function in comparison to *Xdh^fl/fl^*mice were observed^38,56^. The reason for this discrepancy may relate to the use of a mixed-age population of mice, power of the study and to the sex utilised in the Kelley study. In the aforementioned study, mice were 6-13 weeks old, with an n-number (n=7) and only male mice were used. Our study used equal numbers of male and female mice and aged-matched littermates, as well as assessment of a time-course exposing important age-dependent changes^57^.

Mechanistically, our findings link the XOR-dependent vascular dysfunction temporally with inflammation. Within the liver, heart and plasma we saw consistent elevations in one of the early cytokine signals in the inflammatory response i.e. TNFα. These differences overall appeared to increase with age, particularly in the HXOR KO, but also appeared to follow a temporal pattern that begins in early age with rises in the cytokine in the liver, then plasma and then the heart. We suggest that the removal of *Xdh* from the liver eliminates an NO-mediated brake-mechanism leading to a proinflammatory phenotype within the liver and release of TNFα into the systemic circulation causing the consequent endothelial dysfunction and cardiac phenotype that we observed.

We also demonstrated an increased albumin-bilirubin (ALBI) score, indicative of liver dysfunction. Impaired liver function affects the synthesis and metabolism of vasoactive substances, disrupts bile acid regulation, and contributes to the dysregulation of systemic hemodynamics. An increased ALBI score is associated with increased mortality risk in heart failure patients^58^. Whilst HXOR KO mice did not express increased levels of traditional markers of hepatocyte damage (ALT, AST, and ALP) this could be attributed to the collection of plasma used for the biochemical analysis interfering with the sensitivity of the assays^59^. However, ALBI score has demonstrated an ability to show earlier indications of liver disease than the traditional prognostic biomarkers^60^. These observations do highlight, however, the impact of liver health upon cardiac and vascular function.

We sought to investigate a physiological role for XOR mediated bioconversion of nitrite to NO in the setting of leukocyte recruitment, since a key property of endothelium-derived NO is its anti-inflammatory effect^61^. Under basal conditions leukocyte rolling and adhesion were elevated in *Xdh^+/-^* mice compared to *Xdh^+/+^*, with the same magnitude of response observed in HXOR KO mice vs *Xdh^fl/fl^*. This is in contrast to a study in cats treated with allopurinol (under basal conditions) where no change in leukocyte adhesion or emigration was evidenced compared to control^62^. This discrepancy can likely be explained by the fact that in this study cats were administered heparin before assessment. Heparin will compete with endothelial GAGs, to which circulating XOR binds, and thus reduce surface endothelium XOR expression and therefore impacting leukocyte binding^63^. We suggest that there is a dual action of endothelial GAG-bound XOR: purine degradation resulting in superoxide and hydrogen peroxide production both of which elicit leukocyte recruitment^62^, and nitrite reduction to NO which inhibits leukocyte recruitment^64,65^. Our observation suggests that under basal conditions, the reduction in NO production has a more prominent effect on leukocyte recruitment than the reduction in ROS generation.

It is well accepted that NO reduces leukocyte endothelium interactions and hence recruitment. The mechanisms by which it does this include suppression of CD62P, CD62E and VCAM-1 expression on endothelial cells^66,67^. Indeed, in *Xdh^+/-^* and HXOR KO mice we observed an increase in endothelial CD62P expression within the liver, which likely accounts for the observed elevated leukocyte rolling, in line with previous data demonstrating that NO regulates endothelial CD62P expression^68^. These results suggest that under basal conditions, hepatic XOR likely binding to endothelium via interaction with GAGs leads to XOR-dependent nitrite derived NO that plays a role in sustaining a quiescent endothelium. Since XOR is implicated in I/R injury (for review see^69^) an effect attributed to XOR-derived ROS^62^, and oxidative stress-induced expression of TNFα and MCP-1^70^ we assessed the impact of XOR deletion in a mesenteric artery I/R injury model. We observed an increase in leukocyte recruitment in *Xdh^+/+^* mice subjected to I/R injury, with no change in *Xdh^+/-^* mice; an effect concordant with observations by others assessing the impact of renal I/R injury in a similar mouse^70^. Collectively our observations confirm that reducing XOR-derived ROS in the setting of I/R injury is beneficial. But that under physiological conditions XOR-derived NO is important for maintaining vessel homeostasis to limit leukocyte recruitment and vascular dysfunction. There are limited studies investigating XOR expression on leukocytes however we did identify XOR expression in neutrophils (others have suggested XOR expression in macrophages predominantly^71,72^). However, we saw no change in activation state of any leukocyte subtype assessed, supporting our contention that the anti-leukocyte effects observed likely relate to endothelium-bound XOR.

A complication of XOR inhibition *in vivo* is that using allopurinol/febuxostat or genetic deletion, will inhibit UA, ROS and NO production making identification of which pathways are the key drivers for any phenotype seen complex. An alternative method for targeting XOR is by providing an excess of substrate that would drive one of the synthesis pathways. Indeed, to assess XOR-dependent nitrite reduction comparison of the effects of elevating circulating nitrite as a substrate provides such an approach. In our study, dietary inorganic nitrate treatment elevated circulating nitrite in wild types and XOR-deleted mice. This rise in nitrite was associated with a reduction in leukocyte rolling in *Xdh^+/+^*mice subjected to I/R injury, whilst there was no effect in *Xdh^+/-^*, confirming XOR as a nitrite reductase and demonstrating a functional effect *in vivo*.

In summary, we have demonstrated a critical role for the nitrite reductase activity of hepatic XOR *in vivo*. Our data support the hypothesis that in physiological conditions hepatic XOR functions as a nitrite reductase to sustain cardiovascular health. Additionally, in diseases such as hypertension, heart failure or coronary artery disease, conditions associated with reductions in circulating nitrite and nitrate, that correcting circulating nitrite levels using inorganic nitrate to enhance XOR activity, and thus NO generation, might be a preferable therapeutic strategy (as shown recently^73,74^) rather than inhibition of the enzyme to reduce UA or ROS. We suggest that these findings highlight the potential for exploitation of XOR as a nitrite reductase, through nitrate/nitrite delivery, in sustaining cardiovascular health and in treating disease where XOR expression is elevated.

## Supporting information

Supplemental Methods and Results

## Acknowledgements

We would like to extend our sincere thanks to the Cortese-Krott lab for their invaluable assistance in establishing their refined FMD protocol, which was instrumental in our study. Their expertise and support significantly contributed to the success of our research.

## CreDit Nomenclature

ND: data collection, data curation, formal analysis, data interpretation, writing – original draft and takes responsibility for the data.

RSK: data collection, data curation, formal analysis, data interpretation, writing – original draft

TP: data collection, reviewing– original draft

GM: data collection, reviewing– original draft

NHHK: data collection, reviewing– original draft

AAN: data collection, reviewing– original draft

LCG: data collection, reviewing– original draft

IL: data collection, reviewing– original draft

US: data collection, reviewing– original draft

AJS: data collection, reviewing– original draft

JWH: data collection, reviewing– original draft

KSR: data collection, reviewing– original draft

MRB: formal analysis, reviewing– original draft

CPC: formal analysis, reviewing– original draft

AA: conceptualisation, funding acquisition, investigation, methodology, data interpretation, writing – original draft and takes responsibility for the data.

## Data Availability

The data that support the findings of this study are available on reasonable request from the corresponding author, [AA].

## Supplemental Material

Supplemental Methods

Tables S1–S9

Figure S1-S16

References #1–#2

