## Supplemental Methods and Results for "The nitrite reductase activity of xanthine oxidoreductase sustains cardiovascular health"

**Short title:** XOR NO_2_^-^ Reductase Activity in Cardiovascular Health

^+^To whom correspondence should be addressed:

Amrita Ahluwalia, Barts and The London Faculty of Medicine & Dentistry, Queen Mary University of London, Charterhouse Square, London EC1M 6BQ, United Kingdom.

### Supplementary materials and methods

#### General husbandry

All mice were housed in individually ventilated cages with tubes for environmental enrichment included and access to food and water ad libitum. A total of Xdh*^+/+^* = 114, *Xdh^+/-^* = 132, *Xdh^-/-^* = 30 mice and *Xdh^fl/fl^* = 91, HXOR KO = 93 were used.

#### Generation and identification of *Xdh* knockout mice­­­­­­­

Briefly, portions of exon 3 and exon 4 and the entire intron 3 were replaced by a phosphoglycerate kinase/neo cassette to disrupt the *Xdh* gene. Embryonic stem cells were deposited with the mutant mouse resource and research centers (MMRRC). The embyros were used to generate wild type (*Xdh^+/+^*) and heterozygote (*Xdh^+/-^*) mice by MRC Harwell, UK. *Xdh^+/-^* breeding trios were setup in-house for breeding purposes and resulting *Xdh^+/+^*, *Xdh^+/-^* and *Xdh^-/-^* mice used for studies.

Overt physical characteristics were only present in *Xdh^-/-^* mice which were typically smaller than littermates at 2 weeks of age and mortality readily begins to occur due to renal failure. Dissection of the mice revealed all organs visually normal except for the kidneys which were small and pale/white due to fibrosis.

In order to identify the genotype of the offspring ear clips were obtained and lysed overnight at 55^o^­­C using proteinase K (0.3mg/ml) in DirectPCR ear lysis buffer (Bioquote­, UK). The samples were then incubated at 85^o^C for 45 min after which they were kept at 4^o^C. Two sets of primers were utilised to identify the *Xdh^+/+^* and *Xdh^-/-^* sequence. *Xdh^+/+^* primers consisted of forward 5'-CCTATGCCTTCCACAGTTGT-3' and reverse 3'-CACCGTGATGATCTCCAAGT-5' and *Xdh^-/-^* primers consisted of forward 5'-ATGCGATGTTCGCTTGGTGG-3 and reverse 3'-CTATTCGGCTATGACTGGGC -5'. For each sample 2 master mixes were used containing 2x Biomix Red (Bioline, UK) and either *Xdh^+/+^* or *Xdh^-/-^* primers, which was added to 1µl ear lysed sample. The PCR conditions used were 94°C for 3 min followed by 35 cycles at 94°C for 30 s, 60°C for 30 s, and 72°C for 1 min, followed by 72°C for 7 min. The amplified products (*Xdh^+/+^* 1283bp and *Xdh^-/-^* 350bp) were visualized on a 1.5% agarose gel using a FlourChem Imager (Protein Simple, USA).

#### Generation and identification of *Xdh* – Hepatocyte Specific KO

C57BL/6J *Xdh* floxed mice were generated by MRC Harwell via CRISPR/Cas9 genome editing. LoxP sites were inserted into introns 5 and 6 to flank exon 6 of the *Xdh* gene located on chromosome 17. Integration of LoxP at the target sites was validated using Sanger sequencing. Cre mediated excision of floxed *Xdh* exon 6 (and parts of surrounding introns) results in a frameshift mutation and a premature stop codon after 210 amino acids. C57BL/6J Albumin (Alb) Cre mice (purchased from The Jackson Laboratory) were crossbred with *Xdh* floxed mice (*Xdh^fl/fl^*) to produce mice heterozygous for both Alb Cre and *Xdh* flox. Breeding trios were set up using animals homozygous for floxed *Xdh* i.e. *Xdh^fl/fl^* and heterozygous for Alb Cre expression. Overt distinct physical characteristics were not present in HXOR KO mice compared to their controls.

Two sets of primers were utilised to identify the *Xdh* Flox and Alb Cre expression. *Xdh^fl/lf^* primers consisted of forward 5'-AAGGCAAGCAGACGCTACC-3' and reverse 3'-GCCAAGTCTCCAGGCATTGTA-5' and Alb Cre primers consisted of forward 5'-TGCAAACATCACATGCACAC-3, reverse 3'-TTGGCCCCTTACCATAACTG-5', and mutant 5'-GAAGCAGAAGCTTAGGAAGATGG-3'. *Xdh^fl/fl^* PCR was performed using HotStarTaq Master Mix Kit (Qiagen) and 0.4 μM of forward and reverse primers were added to 0.5 µl ear lysed sample. Alb-Cre PCR was performed using BioMix™ Red (Bioline, UK) using 0.4 μM of forward, reverse, and mutant primers. The PCR conditions used for *Xdh^fl/fl^* were 95°C for 15 min followed by 35 cycles at 95°C for 1 min, 53°C for 1 min, and 72°C for 30 s. The PCR conditions used for Alb Cre were 94°C for 3 min followed by 35 cycles at 94°C for 30 s, 60°C for 30 s, and 72°C for 1 min, followed by 72°C for 2 min. The amplified *Xdh* Flox products (*Xdh^+/+^* 125 bp and *Xdh^fl/f^* 139 bp) and Alb Cre products (WT 350 bp and Alb Cre 391 bp) were visualized on a 1.5% agarose gel using a FlourChem Imager (Protein Simple, USA).

#### Tail cuff blood pressure measurement

Mice were gently placed in a restrainer and left to acclimate in a custom-made heat box (Clinipath Ltd, UK) with an ambient temperature of 30°C for 5 min prior to beginning recordings. Mice were subjected to five acclimation cycles followed by a further ten cycles of data acquisition. Invalid cycles due to animal movements or insufficient blood flow to the tail were automatically discarded by the CODA software. Blood pressure (BP) was measured over three consecutive days for each time point and conducted at the same time throughout the study in a quiet environment with minimal disruption. Each mouse was measured individually. Average mean arterial pressure (MAP), systolic BP (SBP) and diastolic BP (DBP) were calculated to generate a single n for each mouse. Any data points +/- 15 mmHg from the average of the 10 data cycles were excluded from final analysis.

#### Echocardiography

Anaesthesia was induced using 5% Isoflurane with 0.4 L/min oxygen and maintained using 1.5 - 2% Isoflurane with 0.4 L/min oxygen. Redux Electrolyte Crème (Parker Laboratories, Inc., USA) was placed on the back of each paw and the mouse secured in a supine position, on a Vevo mouse-handling platform (FujiFilm VisualSonics Inc., Netherlands), with paws taped to electrodes allowing observation of ECG and breathing rate. Body temperature was monitored using a rectal probe and maintained manually between 36.5 – 37.5°C using a heated stage and a heat lamp. Chest fur was removed using Veet® For Men Hair Removal Gel Cream (available from Boots, UK) and Aqua Gel® Lubricating Gel (CIVCO Medical Solutions, USA) applied liberally to the chest. Three separate images were acquired for each field of view and cardiac parameter in B-mode, M-mode, doppler, and tissue doppler. Analysis was performed blinded, and measurements and calculations were generated in Vevo LAB 5.5.1 software. Three measurements from each parameter were averaged to generate a single n. The data shown represent mean ± SEM for each treatment group.

#### Flow Mediated Dilation (FMD)

Flow mediated dilation of the arteria iliac externa was visualized using a Vevo 3100 imaging system with a MX550D, 40MHz transducer (FujiFilm VisualSonics Inc., Netherlands) and the acquisition was carried out blind. Anesthesia was induced using 5% Isoflurane with 0.4 L/min oxygen and maintained using 1.5 - 2% Isoflurane with 0.4 L/min oxygen. Redux Electrolyte Crème (Parker Laboratories, Inc., USA) was placed on the back of each paw and the mouse secured in a supine position, on a Vevo mouse-handling platform (FujiFilm VisualSonics Inc., Netherlands), with paws taped to electrodes allowing observation of ECG and breathing rate. Body temperature was monitored using a rectal probe and maintained manually between 36.5 – 37.5°C using a heated stage and a heat lamp. Hind left limb and abdomen fur was removed using Veet® For Men Hair Removal Gel Cream (available from Boots, UK) and Aqua Gel® Lubricating Gel (CIVCO Medical Solutions, USA) applied liberally to the chest. An O-cuff (Kent Scientific Corporation, USA) was placed around the hind left limb knee. Baseline B-mode acquisition the arteria iliaca extern was recorded 3 times. Confirmation of the arteria iliaca extern was confirmed via pulsatile blood flow via the pulse wave doppler function. The O-cuff was inflated to 300 mmHg via a KAL 84 Calibration Device (Halstrup-Walcher, Germany) and B mode acquisition of the vessel was conducted at 1-min intervals for 5 min. At the 5-min mark the cuff was rapidly deflated to 0 mmHg and subsequent B mode acquisition of the arteria iliaca was acquired at 20 s intervals for 5 min. Analysis was performed blinded, and 5 measurements of the vessel diameter at diastolic pulse, in the same location for each time point, was averaged and referenced to the fold change of the baseline diameter in Vevo LAB 5.5.1 software. The data shown represent mean ± SEM for each treatment group.

#### Intravital microscopy to determine leukocyte recruitment

Four week old mice were anaesthetised using xylazine 7.5mg/kg and ketamine 150mg/kg i.p., the mesentery exposed and superfused with bicarbonate-buffered solution (BBS) [132 mmol/L NaCl, 4.7 mmol/L KCl, 1.2 mmol/L, MgSO, 17.9 mmol/L NaHCO3, and 2.0 mmol/L CaCl2 (pH 7.4)], gassed with 5% CO_2_ and 95% N_2_ at 37°C. Leukocyte rolling and adhesion was counted three times in three vessels in each animal and these values averaged to represent a single n value. A suitable venule was identified as being 20–40 μm in diameter and 100 μm in length. Leukocyte rolling was counted as the number of leukocytes rolling past a fixed point in 1 min and adhesion was identified as a leukocyte remaining stationary for >30 s in a 1-min period.

For ischaemia-reperfusion studies animals were anaesthetised as above and the superior mesenteric artery was clamped for 30 min and allowed to reperfuse for 45 min, after which leukocyte recruitment was determined in one vessel per animal.

#### Platelet isolation and cGMP Enzymeimmunoassay

20-week-old mice were anaesthetised using 5% isoflurane/ 1.5 L/min O_2_ and maintained with 2.5-3.0% isoflurane/ 1.5 L/min O_2_. Blood was extracted via cardiac puncture using a 23G needle into a 1 ml syringe containing 70 µl 3.8% (w/v) sodium citrate (Sigma Aldrich). 3-Isobutyl-1-methylxanthine (IBMX) stock of 100 mM (Sigma Aldrich®, UK) was prepared in dimethyl sulfoxide (Sigma Aldrich®, UK) and subsequently diluted with sterile saline to 10 mM. 600 μl of the citrated blood was aliquoted and IBMX was added to give a final concentration of 100 μM. An equal volume of a modified Tyrode’s/HEPES (MTH) buffer (5 mM glucose, 1 mg/ml BSA, 134 mM NaCl, 20 mM HEPES, 2.9 mM KCl, 0.34 mM Na_2_HPO_4_, 1 mM MgCl, and 12mM NaHCO_3_) was added to produce a 1:2 diluted blood sample. An initial “soft” centrifugation at 120 *g* at 4 ^o^C for 10 min was conducted. The plasma supernatant and buffy coat were collected and 0.02 U/ml apyrase and 2 μg/ml PGI2 (Cambridge Bioscience Ltd, UK) were added to prevent platelet activation. A second “hard” centrifugation at 1000 *g* at 4 ^o^C for 10 min of the platelet rich plasma, pelleted the platelets. The platelet pellet, supernatant, and RBCs were snap frozen in in liquid nitrogen and stored at - 80 ^o^C.

A commercially available cGMP EIA kit “Amersham cGMP Direct Biotrak™, (Cytiva Life Sciences, USA) was performed according to manufacturer’s instructions. Platelets were lysed in 120 μl “Lysis reagent 1” and 12μl of “acetylation reagent” (1-part acetic anhydride and 2-parts trimethylamine) were added to each sample and vortexed. 100 μl of antiserum was added into all wells, of a precoated microplate (except for a blank and a non-specific binding NSB well) and 50 μl of samples and standards (512-2 fmol cGMP) were loaded in duplicate and then incubated for 2 h at 4°C. After this 100 μl of diluted conjugate was added to all wells (except for the blank) and incubated for 1 hour at 4°C. The contents of the wells were aspirated and washed with 200 μl of wash buffer. This step was repeated 4 times before the addition of 200 μl enzyme substrate to each well and incubated for 30 min at room temperature. 100 μl of 1M sulphuric acid was added to all wells, to stop the reaction, and the optical density was recorded at 450nM wavelength within 30 min (MRT-TC Revelaion, Dynex Technologies, UK). The cGMP EIA is dependent on competitive binding between the unlabelled cGMP in the platelet sample and a fixed quantity of peroxidase labelled cGMP. Therefore, increased TMB mediated optical density, at 450 nm is inversely proportional to platelet cGMP binding.

#### TNF-α ELISA

Plasma TNF-α levels were determined using a commercially available DuoSet ELISA kit composed of the Mouse TNF-α Assay Kit, according to manufacturer’s instructions (DY410-05, R&D Systems, USA).

#### Uric Acid Assay

Uric acid quantification was determined using a commercially available kit and was performed according to manufacturer’s instructions. (Uric Acid Assay Kit MAK077, Sigma-Aldrich). Plasma was diluted 1:4 with the provided uric acid buffer and 50 μL of samples and standards (4.0-0.2 nmol/well) were loaded in duplicate in a 96 well plate. 50 μL of the reaction master mix was added to each well and the plate was incubated in the dark at 37 ^o^C for 30 min. Light absorbance was measured at 570 nm (MRX-TC Revelation, Dynex Technologies) and sample uric acid concentration was determined from the standard curve.

#### Flow cytometry

All antibodies were purchased from eBioscience unless otherwise stated. Blood was collected by cardiac puncture into 3.8% sodium citrate and stored on ice. Two flow cytometry panels were used, the first being anti-mouse Ly-6G (Gr-1) FITC (1:250; RRID:AB_465314), anti-mouse CD115 APC (1:33; RRID:AB_1210790), anti-mouse CD11b eFlour450 (1:60; RRID:AB_1582236), anti-mouse CD62L APC-efluor 780 (1:60;  RRID:AB_1603256), anti-mouse CD162 PE (1:8; BD Biosciences; RRID:AB_395719) and anti-mouse CXCR2 (CD182) PE-VIO 615 (Miltenyi Biotec, RRID:AB_2727136). The second panel used anti-mouse CD3 PE-Cy7 (1:8; RRID:AB_469571), anti-mouse CD4 APC (1:33; RRID:AB_469320), anti-mouse CD8a FITC (1:20; RRID:AB_464915), anti-mouse CD19 Brilliant violet 605 (1:10; Biolegend; RRID:AB_11203538) and also anti-mouse CD11b, CD62L and CD162 described above. 50 μl blood was incubated with the above antibodies or the appropriate isotype controls for 30 minutes at 4^o^C in the dark following which erythrocytes were lysed using RBC lysis buffer (eBioscience, UK).

To detect leukocyte XOR expression, blood was collected by cardiac puncture into 3.8% sodium citrate and mixed 1:1 with PBS. Peripheral blood mononuclear cells (PBMCs) and polymorphonuclear cells (PMNs) were isolated using density centrifugation. Histopaque 1077 (Sigma, UK) was layered on top of histopaque 1119 (Sigma, UK) and the diluted blood layered above, followed by centrifugation at 700 *g*, 30 min, room temperature with the brake off. PBMCs and PMNs were collected and seeded in a 96 well plate for antibody staining. Cell pellets were incubated in 2% human and donkey serum blocking buffer for 15 min and subsequently labelled for GR1, CD115, C19 (as above) and anti-mouse CD3 PE (RRID:AB_312663) and anti-mouse xanthine oxidase (1:1000; ab133268, Abcam, UK; RRID:AB_11154903), for 30min at 4^o^C. Cells were washed and the secondary antibody for XO (donkey anti-rabbit IgG preadsorbed, Abcam, RRID:AB_2715515) incubated for 30 min 4^o^C. Cells were then washed twice, fixed (Intracellular fixation and permeabilisation buffer, Thermofisher) and subsequently stored in PBS before acquisition to assess extracellular expression. In order to assess intracellular expression, cells were permeabilised (Intracellular fixation and permeabilisation buffer, Thermofisher) and the cells incubated in blocking buffer followed by primary XO antibody and secondary antibody as described above.

To determine bone marrow neutrophil and inflammatory monocyte CXCR2 expression, bone marrow cells were isolated from femurs by removing the ends and flushing with ice cold PBS. The flow through was centrifuged at 400 g for 5 min at 4^o^C and the cell pellet collected. The cell pellet was treated the same as blood used for detection of leukocyte activation markers (described above), however cells were blocked with 1:100 Fc receptor block (anti-mouse CD16/32; RRID:AB_467135) before antibody incubation. All flow cytometric data was analysed using FACS Diva software (BD Biosciences).

#### Measurement of nitrate and nitrite levels

Plasma and tissue nitrite and nitrate levels (collectively termed NO_x_) were measured using the ozone-based chemiluminescence method. Blood samples were collected from anesthetized animals by cardiac puncture into 1 mL syringes containing 0.08 mL 3.8% (wt/vol) sodium citrate and centrifuged at 4 °C, 13,000 *g* for 5 min and the plasma collected. Plasma and tissue samples were collected, snap frozen, and stored at −80 °C. Tissue samples were homogenized in the presence of a protease inhibitor mixture containing 4-(2- Aminoethyl)benzenesulfonyl fluoride (1 mg/mL), antipain, aprotinin, benzamidine, leupeptin, and pepstatin A, all at a concentration of 10 μg/mL, using a Precellys homogenizer at 4 °C and the homogenate centrifuged at 10,000*g*, 5 min, 4 °C, and the supernatant collected. Plasma and tissue supernatant were filtered using Sartorius Vivaspin 500 3,000 molecular weight cut-off PES (Sartorius Stedim Biotech) at 4 °C, 14,000 *g* for 60 min (plasma) or 90 min (tissue). Before use, filters were washed twice with low NOx containing 18 MΩ dH_2_O. To determine total NO_x_ concentration, samples were added to 0.1 mol/L vanadium (III) chloride in 1 mol/L hydrochloric acid refluxing at 95 °C under N_2_. Nitrite concentration was determined by addition of samples to 0.09 mol/L potassium iodide in glacial acetic acid under nitrogen at room temperature. Nitrate concentration was calculated by subtraction of the nitrite concentration from the total NO_x_.

#### Immunohistochemistry

Animals were perfused with saline via the left ventricle followed by 10% formal saline at 100 mmHg (physiological pressure) for 2 min. The heart, liver and aorta were excised and stored in formal saline overnight and the following day transferred to sterile saline for subsequent analysis. Some liver samples were frozen whilst all other tissue samples were paraffin wax embedded and sectioned. Heart and liver samples were stained with H&E, CD62P (1:100; RRID:AB_2285644) and CD45 (1:2000; RRID:AB_442810) and picrosirius red. Images were acquired using Panoramic 250HT (3D Histech) and analysed using ImageJ software (NIH). For CD62P analysis, 6 vessels per section (20-40 μm) were randomly selected and the area of staining quantified, which was normalised to the circumference of each vessel. For CD45 analysis six vessels per section (20-40 μm) were randomly selected and a region of interest (140 μm x 140 μm) drawn around each vessel and the number of CD45+ cells counted. To determine perivascular collagen deposition, picrosirius red staining was quantified around four randomly selected vessels, and normalised to the circumference of each vessel. For interstitial collagen, four images containing no vessels were acquired and quantified.

Wheatgerm agglutinin fluorescent staining was conducted in order to determine cardiac myocyte area. Sections were stained with wheat germ agglutinin conjugated to Alexaflour 647 (1:500 Molecular Probes, Invitrogen W32466) and Prolong gold DAPI mountant. Images were acquired using a NanoZoomer slide scanner (Hamamatsu) and viewed using NDP software. Four images were taken, and the cardiac myocyte area was analysed using ImageJ, with a minimum of 300 cells analysed per sample. All histological analysis was conducted blinded.

#### Western blotting

Tissue was incubated in stock solution of tissue lysis buffer (10 mmol/L Tris·HCl, 50 mmol/L NaCl, 30 mmol/L NaPPi, and 2 mmol/L EDTA) and 0.5 mol/L NaF, 1% Triton X-100, 0.2 mol/L Na_3_VO_4_, as well as 1 μg/mL each of the protease inhibitors benzamidine, aprotinin, antipain, leupeptin, pepstatin A, and AEBSF. Equal amounts of protein were subjected to 10% (wt/vol) SDS gel electrophoresis under reducing conditions. Separated proteins were then electrotransferred onto 0.2-μm nitrocellulose membrane (GE Healthcare) using semidry electrophoretic transfer cell. Blots were blocked with 5% (wt/vol) BSA and incubated overnight at 4 °C with primary antibody XOR; rabbit polyclonal anti-XOR antibody (XOR 1:2,000; Abcam Cat No 133268; RRID:AB_11154903), primary antibody (p)-eNOS; rabbit polyclonal anti-p-eNOS s1177 antibody ((p)-eNOS 1:1000; Cell Signalling Cat No mAb #9570 or primary antibody total-eNOS; rabbit polyclonal anti-total-eNOS antibody (eNOS 1:500; Santa Cruz Signalling Cat No Sc-650 ) overnight. Membranes were then incubated with goat polyclonal anti-rabbit secondary antibody (1:5,000; Dako) for 1 h and visualized using the ECL Western blotting detection system (FluorChem E Protein Simple). The levels of protein were expressed relative to β‐actin expression (1:10,000; Millipore).

#### Quantitative reverse transcriptase-polymerase chain reaction (qRT-PCR)

All tissue samples were homogenised using a Precellys®24 homogeniser (Bertin Technologies) homogeniser attached to a Cryolys® machine (Bertin Technologies, France) to maintain sample temperatures of 4°C during homogenisation. For liver tissue RNA was extracted, using Nucleospin RNA extraction kit (Macherey Nagel, Germany), according to manufacturer guidelines. Frozen whole hearts were crushed into a homogenous powder via a pestle and mortar and RNA was extracted using RNeasy Fibrous Tissue Kit (Qiagen, UK) according to manufacturer guidelines. cDNA was synthesised and qPCR analysis carried out with SYBR green (ThermoFisher Scientific, UK), using specifically designed primers. See Table S1 for details. qRT-PCR was performed using a QuantStudio™ 7 Flex Real-Time PCR System (Applied Biosystems). Expression was normalised to β-actin and 18s (an average of these two housekeeping genes was used) and expressed as a relative value using the comparative threshold cycle (Ct) method (2−ΔΔCt).

#### RNA sequencing analysis

Quality assessment of the sequencing reads of the 12 samples (6 *Xdh^+/+^* and 6 *Xdh^+/-^* FastQ files) was performed using FastQC (v0.11.9; (http://www.bioinformatics.babraham.ac.uk/projects/fastqc/)) and MultiQC (v1.9; PMID: 27312411). Kallisto (v0.46; PMID: 27043002) was used to index the ensembl *Mus musculus* GRCm39 reference transcriptome and quantify transcripts against the *Mus musculus* GRCm39.104 gtf. Exploratory and differential expression analysis was performed under R (v4.0.5). Transcript abundance was transformed using the rld package from DESeq2 (v2_1.30.1; http://www-huber.embl.de/users/anders/DESeq) for exploratory analysis. Principal component analysis and clustering were used to investigate possible batch effects. Genes with less than 10 counts across samples were removed from further analysis. Differential expression analysis between *Xdh^+/+^* and *Xdh^+/-^* samples was performed with DESeq2 fitting date of birth as a covariate (design ~ DOBL + Genotype). Contrast between *Xdh^+/+^* and *Xdh^+/-^* were extracted and further investigated. All differentially expressed genes with q<0.1 were visualized using the Complex Heatmap package^1^. Ingenuity Pathway Analysis (IPA; QIAGEN;^2^) was used to explore interactions between differentially expressed genes and key biological pathways.

#### Pterin-based fluorometric assay of XDH/XO activity

Undiluted mouse plasma or liver homogenate (prepared as described above) was mixed with pre-warmed PBS (50mM) + Na_2_-EDTA (0.1 mM) buffer pH 7.4 in a final volume of 200μl. Baseline fluorescence was read every 20 s for 5 min as were all subsequent recordings, following drug administration. XO activity was measured by adding 10μM pterin whereas XDH + XO activity was quantified in the presence of 10μM methylene blue. The reaction was inhibited with the addition of 10μM allopurinol and subsequently 1μM isoxanthopterin was added, which is used as an internal standard. Stock solutions were made fresh every day; 100mM pterin and 10mM isoxanthopterin stock solutions were prepared in 1M NaOH and serially diluted. 10mM methylene blue was prepared in MilliQ H_2_O and diluted as required. Fluorescence was measured using a Tecan i-control plate reader excited at 345nm and with emission at 390nm. The fluorescence data was corrected for the immediate fluorescence increase after isoxanthopterin addition and calculated using the following formula:

$$U= \left\{ \Delta F \times\left( \frac{\mathrm{IXPT}}{F_{\mathrm{IXPT}}} \right) \right\}\times0.001 \times\left\{ \frac{V_{c}}{\left( V_{S} \times T \right)} \right\}$$

U is the enzyme activity in µmol/ g tissue / min (plasma is µmol / µl / min), ∆F is the change per second in fluorescence intensity obtained for pterin or methylene blue, [IXPT] is the concentration of the isoxanthopterin added at the end of the assay. F_IPXT_ is the immediate increase of fluorescence obtained after isoxanthopterin addition. V_c_ is the final volume used in mL, V_s_ is the volume of sample added in mL, and T indicate the concentration of the homogenate (mg/ml). The correction factor 0.001 was only used for liver homogenate as plasma did not contain haemoglobin.

### Supplementary tables

| Target Gene | Forward Primer | Reverse Primer |
| --- | --- | --- |
| 18s | AGCCTGCGGCTTAATTTGAC | CAACTAAGAACGGCCATGCA |
| B-Actin | GAAATCGTGCGTGAATCAAAG | TGTAGTTTCATGGAGCCACAG |
| Xdh | GGTTGTTTCCACTTCCTCCA | TTCCAAGGAAACCTCTGTCG |
| eNOS | TCCGGAAGGCGTTTGATC | GCCAAATGTGCTGGTCACC |
| AOX1 | GCCATCTTGTCTGTGCTGTG | CCAGCTTCCGTTCTGACTTG |
| a1-GC | GTCATCACGATGCTCAACGC | GGGTGTCACTCTCTCTGTGC |
| Nox-2 | TTGGGTCAGCACTGGCTCTG | TGGCGGTGTGCAGTGCTATC |
| Nox-4 | GGATCACAGAAGGTCCCTAGCAG | GCGGCTACATGCACACCTGAGAA |
| BNP | TATCTGTCACCGCTGGGAGG | TTGTGAGGCCTTGGTCCTTC |
| TGF-β | GGATACCAACTATTGCTTCAGCTCC | AGGCTCCAAATATAGGGGCAGGGTC |
| SMAD-2 | ATGTCGTCCATCTTGCCATTC | AACCGTCCTGTTTTCTTTAGCTT |
| SMAD-3 | ATGTCAACAGGAATGCAGCAGTGG | ATAGCGCTGGTTACAGTTGGGAGA |
| CTGF | CACAGAGTGGAGCGCCTGTTC | GATGCACTTTTTGCCCTTCTTAATG |
| Fibronectin | CCGTTCCCACTGCTGATTTATC | CCGGTGGCTGTCAGTCAGA |
| Col-1a | TCCTGCTGGTGAGAAAGGAT | TCCAGCAATACCCTGAGGTC |
| NRLP-1 | ATGTGGACCCAACCTTCAAA | GTACGTGCTCCTGGAAAGGT |
| TNF-A | CTGTAGCCCACGTCGTAGC | TTGAGATCCATGCCGTTG |
| IL-1B | GAAATGCCACCTTTTGACAGT | CTGGATGCTCTCATCAGGACA |
| IL-6 | CTGCAAGAGACTTCCATCCAGTT | GAAGTAGGGAAGGCCGTGG |
| IFN-y | AAAGAGATAATCTGGCTCTGC | GTCCTGAGACAATGAACGCT |
| NLRP-1 | ATGTGGACCCAACCTTCAAA | GTACGTGCTCCTGGAAAGGT |
| NLRP-3 | TGCTCTTCACTGCTACTATCAAGCCCT | ACAAGCCTTTGCTCCAGACCCTAT |
| CCL2 | TTAAAAACCTGGATCGGAACCAA | GCATTAGCTTCAGATTTACGGGT |
| CXCL12 | GAGCCAACGTCAAGCATCTG | CGGGTCAATGCACACTTGTC |
| TLR-5 | TTGCTCAAACACCTGGATGC | TCTGCTCACTAGACACACCG |

**Table S1 –** **Primer sequences used for RT-QPCR.** Primer sequences used to determine mRNA expression of genes implicated in canonical/non-canonical NO pathways: xanthine oxidoreductase (*Xdh*) endothelial nitric oxide synthase (eNOS), aldehyde oxidase (AOX1), and soluble guanylyl cyclase α1 (α1-GC). ROS signalling pathways: NADPH oxidase 2 (Nox-2), and NADPH oxidase 4 (Nox-4). Cardiac hypertrophy/heart failure signalling: brain natriuretic peptide (BNP), transforming growth factor β (TGFβ), fibronectin, type I collagen (Col-1a), NLR family pyrin domain containing 1 (NLRP1), tumour necrosis factor α (TNF-α), interleukin 6 (IL-6), and interleukin 10 (IL-10). Two reference housekeeping genes were used as internal controls: ribosomal RNA (18S), and β-actin.

| **Plasma** | ***Xdh^+/+^*** | ***Xdh^+/-^*** | ***Xdh^fl//fl^*** | ***HXOR KO*** | **P-value**  **(*Xdh^+/+^ vs Xdh^+/-^)*** | **P-value**  **(Xdh^fl/fl^ vs HXOR KO)** |
| --- | --- | --- | --- | --- | --- | --- |
| Sodium (mmol/L) | 151.0 ± 3.0 | 154.0 ± 2.3 | 186.5 ± 8.1 | 197.2 ± 8.8 | 0.446 | 0.382 |
| Potassium (mmol/L) | 4.26 ± 0.22 | 4.59 ± 0.26 | 3.63 ± 0.22 | 3.98 ± 0.26 | 0.359 | 0.267 |
| Chloride (mmol/L) | 106.8 ± 0.9 | 107.0 ± 1.4 | 69.10 ± 4.1 | 98.40 ± 5.0 | 0.907 | 0.727 |
| Calcium (mmol/L) | 2.35 ± 0.05 | 2.27 ± 0.02 | 1.42 ± 0.05 | 1.29 ± 0.05 | 0.162 | 0.093 |
| Magnesium (μmol/L) | 0.85 ± 0.03 | 0.80 ± 0.02 | 0.70 ± 0.02 | 0.66 ± 0.02 | 0.148 | 0.092 |
| Urea (mmol/L) | 7.54 ± 0.63 | 5.58 ± 0.42 | 6.73 ± 0.49 | 6.93 ± 0.38 | **0.033** | 0.751 |
| Creatinine (μmol/L) | 9.74 ± 0.88 | 8.92 ± 0.47 | 10.96 ± 0.55 | 11.82 ± 0.69 | 0.433 | 0.343 |
| Total Bilirubin (μmol/L) | 1.68 ± 0.27 | 1.24 ± 0.13 | 1.14 ± 0.08 | 1.48 ± 0.12 | 0.187 | **0.024** |
| Albumin (g/L) | **N/A** | **N/A** | 21.92 ± 0.34 **(9)** | 20.77 ± 0.43 | **N/A** | 0.052 |
| Alkaline phosphatase (ALP; U/I) | 224.8 ± 12.7 | 221.6 ± 9.3 | 48.00 ± 3.6 | 46.70 ± 3.8 | 0.844 | 0.806 |
| Alanine aminotransferase (ALT; U/I) | 18.4 ± 2.5 | 18.0 ± 3.0 | 17.60 ± 1.7 | 15.44 ± 1.1 **(9)** | 0.922 | 0.320 |
| Aspartate aminotransferase (AST; U/I) | 76.2 ± 9.9 | 73.8 ± 16.5 | 74.60 ± 9.6 | 106.0 ± 16.0 | 0.904 | 0.109 |
| Total Cholesterol (mmol/L) | 3.10 ± 0.19 | 3.30 ± 0.19 | 2.01 ± 0.19 | 2.01 ± 0.19 | 0.497 | 0.996 |
| High density lipoprotein (HDL; mmol/L) | 2.18 ± 0.16 | 2.35 ± 0.12 | 1.47 ± 0.16 | 1.46 ± 0.12 | 0.414 | 0.949 |
| Low density lipoprotein (LDL; mmol/L) | 0.56 ± 0.02 | 0.62 ± 0.03 | 0.46 ± 0.05 | 0.44 ± 0.04 | 0.119 | 0.774 |
| Triglyceride (mmol/L) | 1.18 ± 0.14 | 0.96 ± 0.05 | 0.60 ± 0.06 | 0.60 ± 0.06 | 0.171 | 0.922 |
| Free Fatty Acids (mmol/L) | **N/A** | **N/A** | 0.27 ± 0.02 | 0.29 ± 0.02 | **N/A** | 0.337 |
| Creatine kinase (CK; U/I) | 307.2 ± 39 | 301.2 ± 94 | 329.5 ± 78 | 414.6 ± 61 | 0.954 | 0.402 |
| Glucose (mmol/L) | 16.9 ± 0.5 | 16.6 ± 0.3 | 11.82 ± 0.48 | 10.77 ± 0.66 | 0.686 | 0.215 |
| Fructosamine (μmol/L) | 203.0 ± 9.6 | 223.6 ± 6.4 | Not Acquired | Not Acquired | 0.112 | **N/A** |

**Table S2** – Plasma biochemical analysis in 4-week-old *Xdh^+/+^ or Xdh^+/-^* mice and 20-week-old Xdh^fl/fl^ or HXOR KO mice. Unless otherwise stated n=10. Plasma data was analysed for statistical significance using an unpaired Student’s t-test. (1 *Xdh^fl/fl^* and 1 HXOR KO data points excluded using ROUT’s outlier test).

| **Urine** | ***Xdh^+/+^*** | ***Xdh^+/-^*** | ***Xdh^-/-^*** | **P-value** |
| --- | --- | --- | --- | --- |
| Sodium (mmol/L) | 130.2 ± 31.4 | 239.2 ± 57 | 29.4 ± 3.4 | 0.007 |
| Potassium (mmol/L) | 320.2 ± 44.0 | 220.2 ± 46.0 | 44.4 ± 5.6*** | 0.0007 |
| Chloride (mmol/L) | 145.0 ± 22.7 | 211.0 ± 50.4 | 42.8 ± 4.5 | 0.010 |
| Urea (mmol/L) | 1044 ± 62 | 1191 ± 181 | 185 ± 18 (4)*** | 0.0003 |
| Creatinine (μmol/L) | 5376 ± 947 | 4213 ± 511 | 781 ± 88 (4) ** | 0.002 |
| Uric acid (μmol/L) | 1141 ± 269 | 782 ± 139 | 0** ^#^ | 0.002 |
| Urinary protein (mg/dL) | 1313 ± 332 | 546 ± 261.3 | 32.7 ± 13.2 (3) * | 0.037 |

**Table S3** – **Urine biochemical analysis in 4-week-old *Xdh* transgenic mice.** Unless otherwise stated n=5. Plasma *Xdh^-/-^* values were not obtained due to lack of available sample. Urine data was analysed using Two-way ANOVA followed by Dunnett’s post-tests. *P<0.05, **P<0.01, ***P<0.001. ^#^ below detection limit of 89 μmol/L. (Uneven n numbers *Xdh^-/-^* due to lack of sample availability for the required test)

|  | ***Xdh^+/+^*** | ***Xdh^+/-^*** | **P-Value** |
| --- | --- | --- | --- |
| Neutrophil (10^4^ cells/ml) | 8.1 ± 1.9 | 7.5 ± 1.9 | 0.84 |
| Resident monocyte (10^4^ cells/ml) | 5.1 ± 1.7 | 3.6 ± 1.3 | 0.49 |
| Inflammatory monocyte (10^4^ cells/ml) | 0.7 ± 0.3 | 0.7 ± 0.3 | 0.99 |
| T cell (CD3^+^CD4^+^; 10^4^ cells/ml) | 17.8 ± 2.5 | 16.1 ± 2.5 | 0.63 |
| T cell (CD3^+^CD8^+^; 10^4^ cells/ml) | 8.1 ± 1.3 | 7.3 ± 1.4 | 0.67 |
| B cell (CD19^+^; 10^4^ cells/ml) | 45.8 ± 8.6 | 32.7 ± 5.1 | 0.20 |

**Table S4** – **Circulating leukocyte numbers are unaltered in *Xdh^+/-^* mice**. n=12. Statistical significance determined using unpaired Students t test.

|  | ***Xdh^+/+^*** | ***Xdh^+/-^*** | **P-Value** |
| --- | --- | --- | --- |
| **CD162** |  |  |  |
| Neutrophil; Gr1^+^ | 64191 ± 8454 | 79774 ± 7331 | 0.178 |
| RM; CD115^+^ | 49113 ± 6137 | 62082 ± 6193 | 0.151 |
| IM; CD115^+^Gr1^+^ | 106788 ± 14090 | 145491 ± 12646 | 0.053 |
| T cell; CD3^+^CD4^+^ | 26991 ± 3847 | 35302 ± 3088 | 0.106 |
| T cell; CD3^+^CD8^+^ | 42995 ± 5912 | 55729 ± 5488 | 0.110 |
| B cell; CD19^+^ | 1550 ± 206 | 2108 ± 153 | **0.041** |
| % neutrophil; Gr1^+^ expressing | 99.8 ± 0.1 | 99.8 ± 0.1 | 0.653 |
| % RM; CD115^+^ expressing | 99.1 ± 0.3 | 98.3 ± 0.4 | 0.149 |
| % IM; CD115^+^Gr1^+^ expressing | 99.6 ± 0.2 | 98.8 ± 0.5 | **0.067** |
| % CD4^+^ cells expressing | 99.9 ± 0.0 | 99.8 ± 0.0 | 0.041 |
| % CD8^+^ cells expressing | 99.8 ± 0.1 | 99.6 ± 0.1 | 0.166 |
| % CD19^+^ cells expressing | 72.7 ± 6.9 | 76.8 ± 5.4 | 0.648 |
| **CD62L** |  |  |  |
| Neutrophil; Gr1^+^ | 5190 ± 454 | 3838 ± 345 | **0.027** |
| RM; CD115^+^ | 2537 ± 346 | 1877 ± 286 | 0.156 |
| IM; CD115^+^Gr1^+^ | 5453 ± 707 | 4259 ± 554 | 0.197 |
| T cell; CD3^+^CD4^+^ | 2422 ± 249 | 2394 ± 237 | 0.936 |
| T cell; CD3^+^CD8^+^ | 4475 ± 430 | 3119 ± 278 | 0.015 |
| B cell; CD19^+^ | 2185 ± 194 | 1680 ± 156 | 0.055 |
| % neutrophil; Gr1^+^ expressing | 99.9 ± 0.0 | 99.9 ± 0.0 | 0.999 |
| % RM; CD115^+^ expressing | 49.9 ± 3.3 | 48.2 ± 3.60 | 0.737 |
| % IM; CD115^+^Gr1 expressing | 100 ± 0 | 98.8 ± 0.5 | **0.032** |
| % CD4^+^ cells expressing | 93.8 ± 0.4 | 94.9 ± 0.6 | 0.133 |
| % CD8^+^ cells expressing | 97.6 ± 0.4 | 98.4 ± 0.2 | 0.075 |
| % CD19^+^ cells expressing | 84.7 ± 1.1 | 82.4 ± 1.5 | 0.238 |
| **CD11b** |  |  |  |
| Neutrophil; Gr1^+^ | 6064 ± 652 | 5717 ± 486 | 0.674 |
| RM; CD115^+^ | 17218 ± 1163 | 18244 ± 1106 | 0.529 |
| IM; CD115^+^Gr1^+^ | 24098 ± 1495 | 23999 ± 1510 | 0.963 |
| T cell; CD3^+^CD4^+^ | 6671 ± 944 | 5714 ± 747 | 0.435 |
| T cell; CD3^+^CD8^+^ | 5606 ± 559 | 4638 ± 496 | 0.209 |
| B cell; CD19^+^ | 1734 ± 216 | 1912 ± 172 | 0.526 |
| % neutrophil; Gr1^+^ expressing | 99.9 ± 0.1 | 99.5 ± 0.2 | 0.189 |
| % RM; CD115^+^ expressing | 98.3 ± 0.4 | 94.4 ± 2.0 | 0.067 |
| % IM; CD115^+^Gr1 expressing | 99.7 ± 0.3 | 99.9 ± 0.1 | 0.604 |
| % CD4^+^ cells expressing | 8.9 ± 1.2 | 6.4 ± 0.8 | 0.113 |
| % CD8^+^ cells expressing | 11.4 ± 1.5 | 8.0 ± 1.6 | 0.143 |
| % CD19^+^ cells expressing | 2.46 ± 0.3 | 8.1 ± 5.4 | 0.313 |

**Table S5** – **Basal levels of leukocyte surface expression of CD62L, CD162 and CD11b**. Values shown are median fluorescence intensity (MFI) and % expression on respective cell types. IM – Inflammatory monocyte; RM – Resident monocyte. n=12. Statistical significance determined using unpaired Students t test and highlighted in bold where considered statistically significant..

|  | ***Xdh^+/+^*** | | **P-Value** | ***Xdh^+/-^*** | | **P-Value** |
| --- | --- | --- | --- | --- | --- | --- |
|  | **KCl** | **KNO_3_** |  | **KCl** | **KNO_3_** |  |
| Neutrophil (10^4^ cells/ml) | 7.7 ± 1.4 (9) | 3.8 ± 0.6 (8) | **0.022** | 8.6 ± 1.3 (9) | 5.6 ± 1.2 (9) | 0.104 |
| Resident monocyte (10^4^ cells/ml) | 4.0 ± 0.9 (9) | 2.5 ± 0.6 (10) | 0.183 | 5.5 ± 1.1 (11) | 4.0 ± 0.8 (11) | 0.265 |
| Inflammatory monocyte (10^4^ cells/ml) | 0.5 ± 0.1 (9) | 0.1 ± 0.04 (8) | **0.010** | 0.8 ± 0.1 (10) | 0.5 ± 0.1 (10) | 0.264 |
| T cell (CD3^+^CD4^+^; 10^4^ cells/ml) | 17.0 ± 2.0 (9) | 15.4 ± 2.3 (10) | 0.606 | 18.0 ± 2.8 (11) | 18.0 ± 2.7 (11) | 0.988 |
| T cell (CD3^+^CD8^+^; 10^4^ cells/ml) | 6.9 ± 0.8 (9) | 6.0 ± 0.9 (10) | 0.479 | 8.5 ± 1.3 (11) | 8.1 ± 1.4 (11) | 0.837 |
| B cell (CD19^+^; 10^4^ cells/ml) | 42.8 ± 6.7 (9) | 35.6 ± 6.8 (10) | 0.465 | 46.3 ± 8.3 (11) | 40.8 ± 8.4 (11) | 0.649 |

**Table S6** – **Inorganic nitrate reduces circulating innate immune cells in *Xdh^+/+^* mice.** Data shown as Mean ± SEM (n). Statistical significance determined using unpaired Students t test. Uneven n numbers represent exclusion of data using ROUTs test.

|  | ***Xdh^+/+^*** | | **P-Value** | ***Xdh^+/-^*** | | **P-Value** |
| --- | --- | --- | --- | --- | --- | --- |
|  | **KCl (8)** | **KNO_3_ (6)** |  | **KCl (9)** | **KNO_3_ (7)** |  |
| **Blood** |  |  |  |  |  |  |
| Neutrophil; Gr1^+^ | 6988 ± 1309 | 7038 ± 1728 | 0.982 | 7150 ± 1395 | 4939 ± 1068 | 0.251 |
| IM; CD115^+^ | 7190 ± 1174 | 6641 ± 1416 | 0.767 | 7324 ± 1297 | 5492 ± 997 | 0.304 |
| % neutrophil; Gr1^+^expressing | 98.0 ± 0.2 | 97.5 ± 1.0 | 0.600 | 96.7 ± 1.2 | 94.8 ± 2.8 | 0.512 |
| % IM; CD115^+^Gr1^+^ expressing | 98.2 ± 0.9 | 98.5 ± 1.0 | 0.801 | 95.5 ± 1.8 | 90.9 ± 4.2 | 0.282 |
| **Bone marrow** | **KCl (5)** | **KNO_3_ (6)** |  | **KCl (6)** | **KNO_3_ (5)** |  |
| Neutrophil; Gr1^+^ | 1944 ± 318 | 2062 ± 357 | 0.814 | 2161 ± 305 | 2481 ± 434 | 0.551 |
| IM; CD115^+^ | 1345 ± 325 | 1664 ± 311 | 0.498 | 1275 ± 295 | 1746 ± 385 | 0.349 |
| % neutrophil; Gr1^+^ expressing | 70.6 ± 6.5 | 77.2 ± 2.7 | 0.343 | 74.4 ± 5.4 | 81.5 ± 2.4 | 0.292 |
| % IM; CD115^+^Gr1^+^ expressing | 66.6 ± 6.0 | 76.3 ± 5.7 | 0.273 | 65.9 ± 5.4 | 75.5 ± 4.8 | 0.227 |

**Table S7** – **CXCR2 expression on innate immune cells unaffected by inorganic nitrate**. Values shown are median fluorescence intensity (MFI) and % expression on respective cell types. IM – Inflammatory monocyte. Statistical significance determined using unpaired Students t test.

|  | ***Xdh^+/+^*** | | **P-Value** | ***Xdh^+/-^*** | | **P-Value** |
| --- | --- | --- | --- | --- | --- | --- |
|  | **KCl (9)** | **KNO_3_ (10)** |  | **KCl (11)** | **KNO_3_ (11)** |  |
| **CD162** |  |  |  |  |  |  |
| Neutrophil; Gr1^+^ | 46133 ± 4646 | 65346 ± 10096 | 0.114 | 50678 ± 5154 | 58109 ± 6061 | 0.361 |
| RM; CD115^+^ | 35491 ± 6591 | 48503 ± 4797 | 0.124 | 45405 ± 5207 | 49863 ± 4154 | 0.511 |
| IM; CD115^+^Gr1^+^ | 74962 ± 10038 | 108875 ± 16896 | 0.112 | 84723 ± 8896 | 99360 ± 10459 | 0.299 |
| T cell; CD3^+^CD4^+^ | 21610 ± 3947 | 30984 ± 5154 | 0.174 | 23432 ± 3936 | 30348 ± 4695 | 0.272 |
| T cell; CD3^+^CD8^+^ | 31522 ± 5776 | 44252 ± 7030 | 0.185 | 33854 ± 5781 | 43116 ± 6646 | 0.306 |
| B cell; CD19^+^ | 1095 ± 190 | 1714 ± 351 | 0.152 | 1212 ± 232 | 1697 ± 317 | 0.231 |
| % neutrophil; Gr1^+^ expressing | 99.8 ± 0.0 | 99.6 ± 0.1 | 0.099 | 99.8 ± 0.0 | 99.7 ± 0.1 | 0.420 |
| % RM; CD115^+^ expressing | 99.4 ± 0.2 | 99.4 ± 0.3 | 0.900 | 99.4 ± 0.2 | 99.7 ± 0.1 | 0.140 |
| % IM; CD115^+^Gr1^+^ expressing | 100 ± 0.0 | 98.7 ± 0.7 | 0.068 | 99.3 ± 0.4 | 99.1 ± 0.5 | 0.759 |
| % CD4^+^ cells expressing | 100.0 ± 0.0 | 99.9 ± 0.0 | 0.033 | 99.9 ± 0.0 | 100.0 ± 0.0 | 0.346 |
| % CD8^+^ cells expressing | 100.0 ± 0.0 | 99.9 ± 0.0 | 0.095 | 99.9 ± 0.1 | 99.8 ± 0.0 | 0.453 |
| % CD19^+^ cells expressing | 84.9 ± 3.1 | 91.1 ± 1.4 | 0.076 | 84.9 ± 2.7 | 88.1 ± 2.1 | 0.364 |
| **CD62L** |  |  |  |  |  |  |
| Neutrophil; Gr1^+^ | 4752 ± 381 | 4136 ± 454 | 0.326 | 3709 ± 613 | 4282 ± 503 | 0.481 |
| RM; CD115^+^ | 2019 ± 109 | 1964 ± 169 | 0.794 | 1859 ± 192 | 2295 ± 282 | 0.215 |
| IM; CD115^+^Gr1^+^ | 4185 ± 314 | 3939 ± 368 | 0.622 | 3822 ± 390 | 4187 ± 448 | 0.545 |
| T cell; CD3^+^CD4^+^ | 2625 ± 271 | 2583 ± 255 | 0.912 | 2438 ± 291 | 2605 ± 291 | 0.689 |
| T cell; CD3^+^CD8^+^ | 3412 ± 323 | 3339 ± 323 | 0.875 | 3100 ± 413 | 3312 ± 372 | 0.707 |
| B cell; CD19^+^ | 1944 ± 156 | 1954 ± 146 | 0.965 | 1845 ± 193 | 1960 ± 173 | 0.663 |
| % neutrophil; Gr1^+^ expressing | 99.8 ± 0.0 | 99.8 ± 0.1 | 0.736 | 98.6 ± 1.2 | 99.8 ± 0.1 | 0.304 |
| % RM; CD115^+^ expressing | 67.1 ± 8.3 | 67.4 ± 9.4 | 0.982 | 74.1 ± 6.3 | 63.8 ± 9.4 | 0.377 |
| % IM; CD115^+^Gr1 expressing | 100 ± 0.0 | 100 ± 0.0 | 1.000 | 99.0 ± 0.4 | 99.7 ± 0.2 | 0.187 |
| % CD4^+^ cells expressing | 92.3 ± 0.8 | 92.7 ± 0.8 | 0.731 | 87.2 ± 5.0 | 93.1 ± 0.5 | 0.250 |
| % CD8^+^ cells expressing | 98.9 ± 0.2 | 99.6 ± 0.2 | 0.032 | 98.2 ± 1.0 | 99.3 ± 0.1 | 0.282 |
| % CD19^+^ cells expressing | 90.7 ± 1.2 | 88.9 ± 1.9 | 0.429 | 90.3 ± 1.5 | 90.1 ± 0.8 | 0.924 |
| **CD11b** |  |  |  |  |  |  |
| Neutrophil; Gr1^+^ | 6888 ± 512 | 7240 ± 578 | 0.657 | 7620 ± 686 | 7027 ± 730 | 0.560 |
| RM; CD115^+^ | 20228 ± 1833 | 17504 ± 810 | 0.177 | 19992 ± 1848 | 16449 ± 934 | 0.103 |
| IM; CD115^+^Gr1^+^ | 25415 ± 2513 | 24879 ± 1882 | 0.865 | 27917 ± 2382 | 23963 ± 1465 | 0.173 |
| T cell; CD3^+^CD4^+^ | 9715 ± 873 | 9557 ± 758 | 0.892 | 10392 ± 894 | 10107 ± 736 | 0.808 |
| T cell; CD3^+^CD8^+^ | 6862 ± 531 | 7944 ± 509 | 0.160 | 7775 ± 806 | 7448 ± 623 | 0.752 |
| B cell; CD19^+^ | 2296 ± 585 | 3387 ± 567 | 0.199 | 3016 ± 537 | 2984 ± 406 | 0.962 |
| % neutrophil; Gr1^+^ expressing | 99.8 ± 0.1 | 99.9 ± 0.1 | 0.951 | 99.9 ± 0.0 | 99.9 ± 0.0 | 0.771 |
| % RM; CD115^+^ expressing | 90.0 ± 3.3 | 93.7 ± 2.5 | 0.373 | 94.2 ± 1.4 | 96.6 ± 1.4 | 0.250 |
| % IM; CD115^+^Gr1 expressing | 99.6 ± 0.3 | 100 ± 0.0 | 0.134 | 99.7 ± 0.2 | 99.9 ± 0.1 | 0.521 |
| % CD4^+^ cells expressing | 9.5 ± 1.2 | 9.2 ± 1 | 0.836 | 11.0 ± 1.0 | 10.5 ± 0.8 | 0.689 |
| % CD8^+^ cells expressing | 11.9 ± 1.4 | 11.3 ± 1.0 | 0.727 | 14.1 ± 1.4 | 11.1 ± 0.9 | 0.078 |
| % CD19^+^ cells expressing | 3.0 ± 0.4 | 2.2 ± 0.2 | 0.129 | 2.9 ± 0.4 | 3.2 ± 0.6 | 0.777 |

**Table S8** – **Inorganic nitrate has no effect on basal leukocyte surface expression of CD162, CD62L and CD11b**. Values shown are MFI and % expression on respective cell types. IM – Inflammatory monocyte; RM – Resident monocyte. Statistical significance determined using unpaired Students t test

|  | **Extracellular** | | | **Permeabilised** | | |
| --- | --- | --- | --- | --- | --- | --- |
|  | ***Xdh^+/+^*** | ***Xdh^+/-^*** | **P-value** | ***Xdh^+/+^*** | ***Xdh^+/-^*** | **P-value** |
| Neutrophil (Gr1^+^) | 2.57 ± 1.45 | 9.4 ± 3.39 | 0.0775 | 16.71 ± 1.73 | 6.61 ± 0.92 | **<0.0001** |
| Monocyte (CD115^+^) | 6.4 ± 3.92 | 13.66 ± 3.84 | 0.1995 | 11.93 ± 3.05 | 6.97 ± 1.38 | 0.1521 |
| Inflammatory monocyte (CD115^+^Gr1^-^) | 17.97 ± 14.21 | 9.88 ± 3.95 | 0.5886 | 9.57 ± 4.36 | 8.98 ± 1.75 | 0.9016 |
| Resident monocyte (CD115^+^Gr1^-^) | 5.77 ± 4.55 | 8.36 ± 3.17 | 0.6451 | 15.75 ± 8.35 | 8.9 ± 2.13 | 0.4352 |
| Lymphocyte (CD3^+^) | 5.41 ± 2.22 | 10.89 ± 4.8 | 0.3117 | 7.91 ± 2.25 | 6.66 ± 1.32 | 0.6360 |
| B cell (CD19^+^) | 4.96 ± 1.8 | 5.37 ± 2.6 | 0.8988 | 6.72 ± 2.31 | 4.67 ± 1.06 | 0.4275 |

**Table S9 – Circulating Leukocyte XOR expression**. Data is expressed as % increase in median fluorescence intensity vs secondary antibody (for xanthine oxidase) alone. Data analysed for statistical significance using an unpaired Student’s t-test.

### Supplementary figures

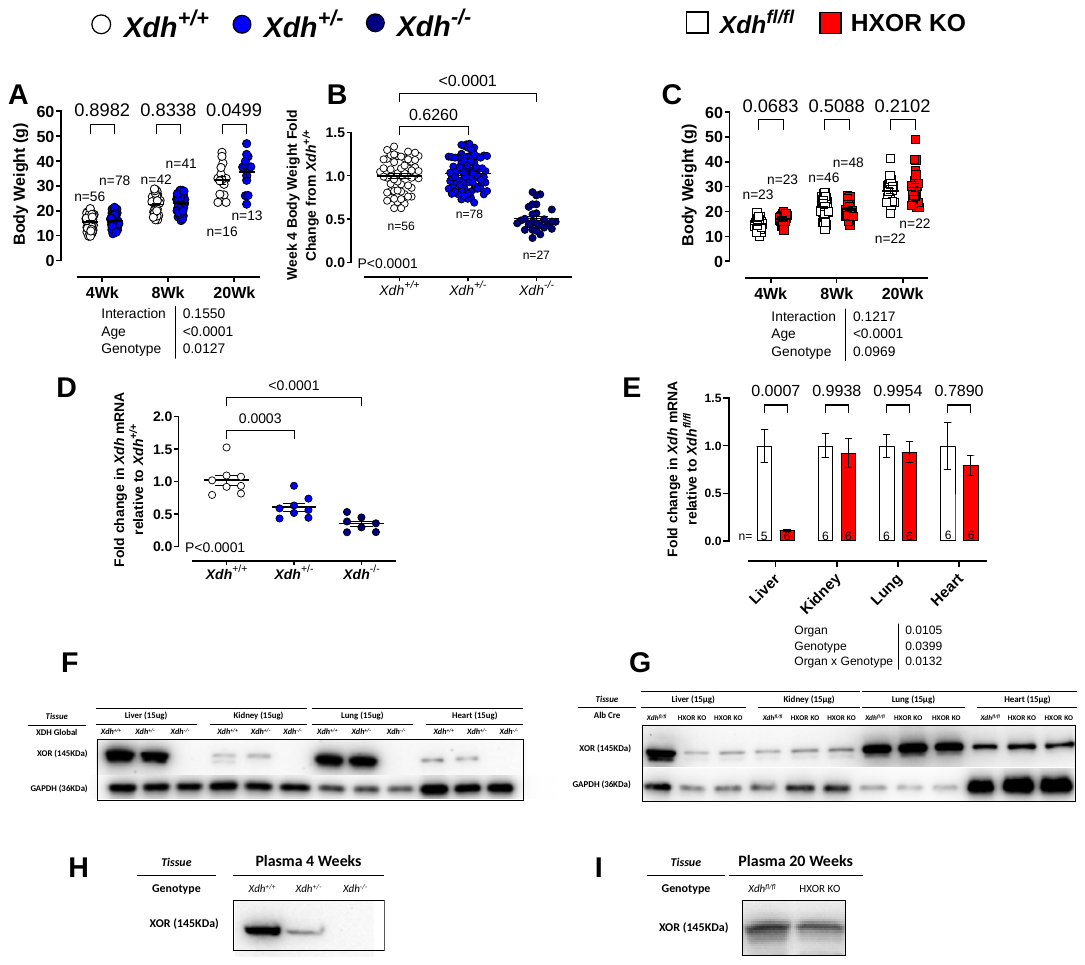
**Figure S1. Confirmation of global and hepatocyte specific XOR KO models. (A)** Body weight of, male and female, *Xdh^+/+^* and *Xdh^+/-^* littermates at 4, 8, and 20-weeks-old. **(B)** fold difference in body weight of, male and female, *Xdh^+/+^*, *Xdh^+/-^*, and *Xdh^-/-^* littermates at 4-weeks-old. **(C)** Body weight of, male and female, *Xdh^fl/fl^* and HXOR KO littermates at 4, 8, and 20-weeks-old. **(D)** Liver *Xdh* mRNA expression in 4-week-old global KO mice. **(E)** *Xdh* mRNA expression in liver, kidney, heart and lungs of 8-week-old HXOR KO mice. Representative Western blot of XOR protein expression in homogenised tissues for each genotype in the **(F)** 4-week-old global KO model and **(G)** 8-week-old HXOR KO model. Representative Western blot of XOR protein expression in the plasma tissues for each genotype in the **(H)** 4-week-old global KO model and **(I)** 20-week-old HXOR KO model. Data are shown as mean ± SEM of n mice (shown on individual graphs). Statistical significance was determined using two-way ANOVA with Sidak’s *post hoc* analysis or using one-way ANOVA followed by Dunnett’s *post hoc* test as appropriate.

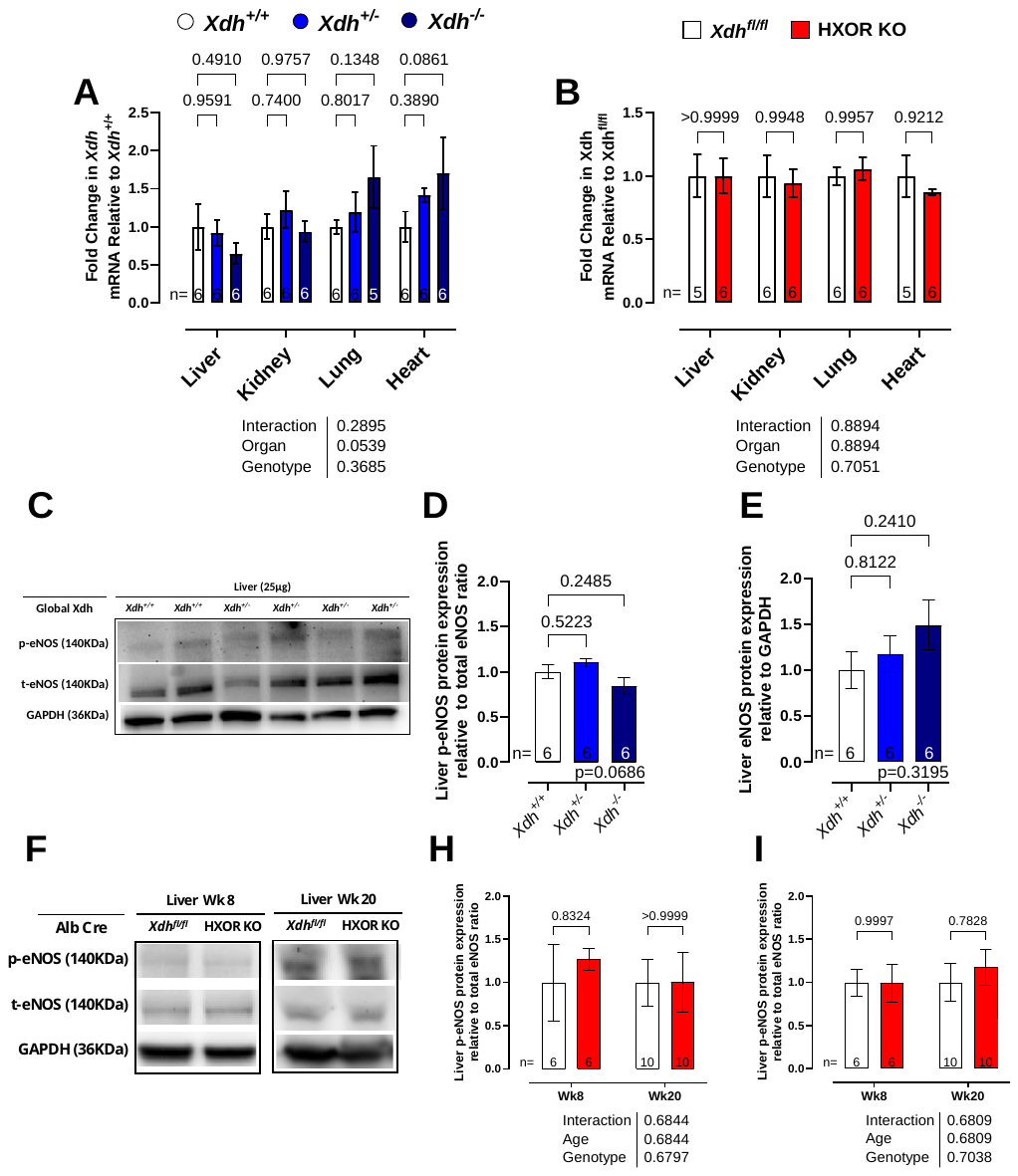
**Figure S2**. Phospho and Total eNOS expression unaffected by removal of *Xdh* eNOS activity and expression in the remains unaltered in HXOR KO mice. (A) NOS3 mRNA expression in the liver, kidney, lungs and heart relative to *Xdh^+/+^* mice, in each tissue assessed, in 4-week-old mice. (B) NOS3 mRNA expression in the liver, kidney, lungs and heart relative to *Xdh^fl/fl^* mice, in each tissue assessed, in 8-week-old mice. (C) Representative Western blot of eNOS phosphorylation and expression in homogenised liver tissue in global *Xdh* KO model mice at 4-week-old. (D) Representative western blot of eNOS phosphorylation and expression in homogenised liver tissue in HXOR KO model mice at 8 and 20-week-old. (D and G) Quantification of Western blots probing phosphorylation of eNOS s1177 relative to total eNOS expression in liver homogenates. (E and H) Quantification of Western blots probing total eNOS expression relative to GAPDH expression in liver homogenates. Data are shown as mean ± SEM of n mice (shown on individual graphs). Statistical significance was determined using two-way ANOVA with Dunnett’s *post hoc* (A, B, G, and H) or One Way ANOVA with Sidak’s *post hoc* analysis (D and E).

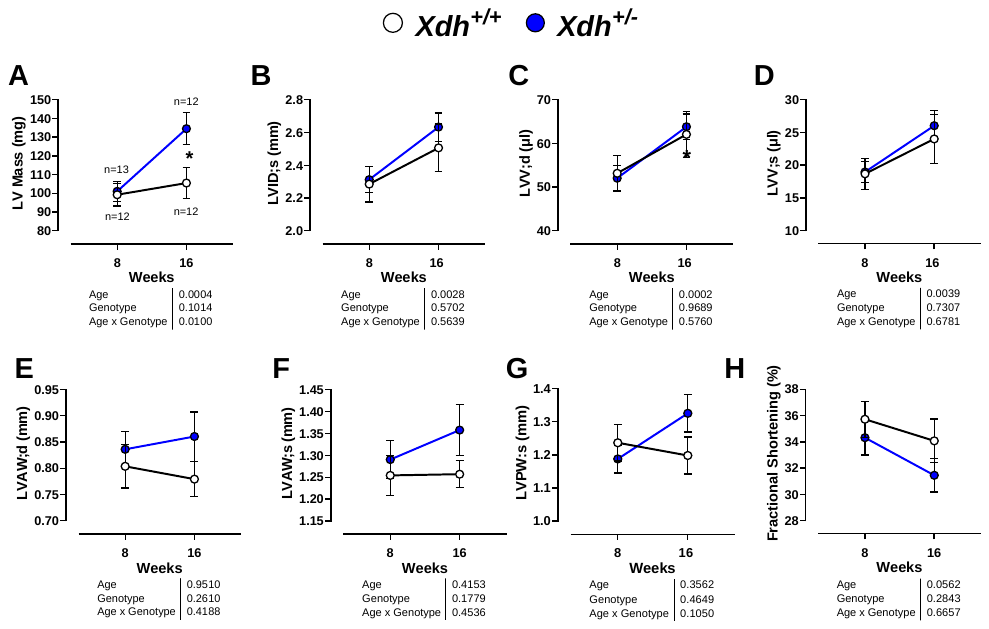
**Figure S3**. *Xdh^+/-^* mice have LV hypertrophy and remodelling. Echocardiography data of *Xdh^+/‑^* versus *Xdh^+/+^* littermates, at 8-16 weeks old. Images were acquired from the LV of mice in short axis view with M-mode measurements for: **(A)** LV mass, **(B)** LV internal diameter: systole, **(C)** LV volume; diastole, **(D)** LV volume; systole, **(E)** LV anterior wall; diastole**,** **(F**) LV anterior wall; systole, **(G)** LV posterior wall; diastole, and **(H)** fractional shortening. Data are shown as mean ± SEM of n mice (shown in **(A)** applies to all graphs). Statistical significance was determined using mixed-effect analysis followed by Sidak’s multiple *post hoc* tests. *Post hoc* test at an individual timepoint (*Xdh^+/+^* vs *Xdh^+/-^*) where relevant is shown as *P<0.05.

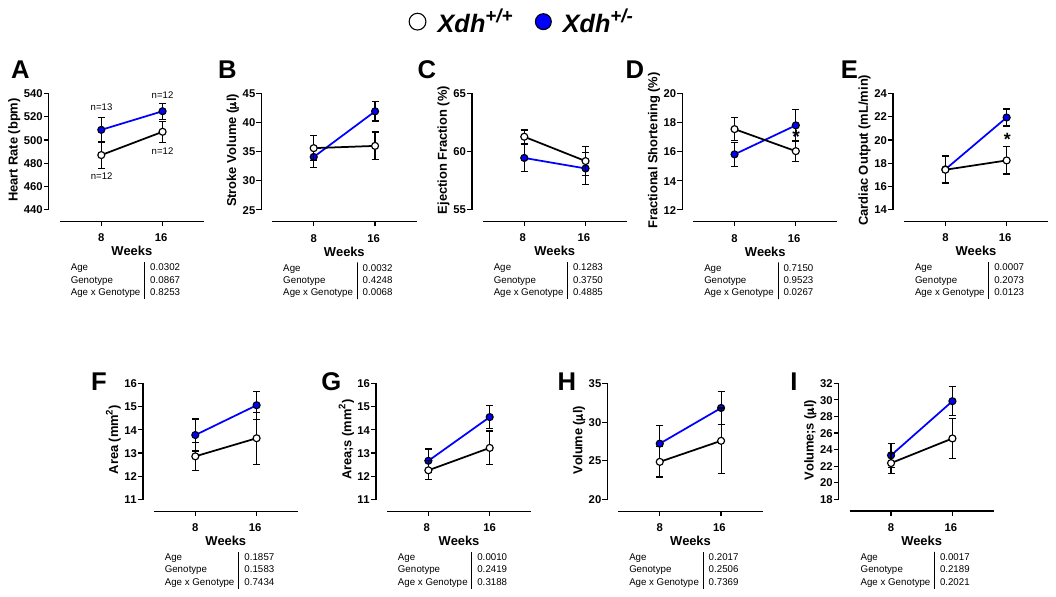
Figure S4 Confirmation of M-mode data that *Xdh^+/-^* mice have LV hypertrophy and remodelling**.** Echocardiography data of *Xdh^+/‑^* versus *Xdh^+/+^* littermates, at 8-16 weeks old Images were acquired from the LV of mice in parasternal long axis view with B-mode measurements: **(A)** heart rate, **(B)** stroke volume, **(C)** ejection fraction, **(D)** fractional shortening, **(E)** cardiac output, **(F)** LV area, **(G)** LV area: systole, **(H)** LV volume, and **(I)** LV volume; systole. Data are shown as mean ± SEM of n mice shown in **(A)** applies to all graphs). Statistical significance was determined using mixed-effect analysis followed by Sidak’s multiple *post hoc* tests.

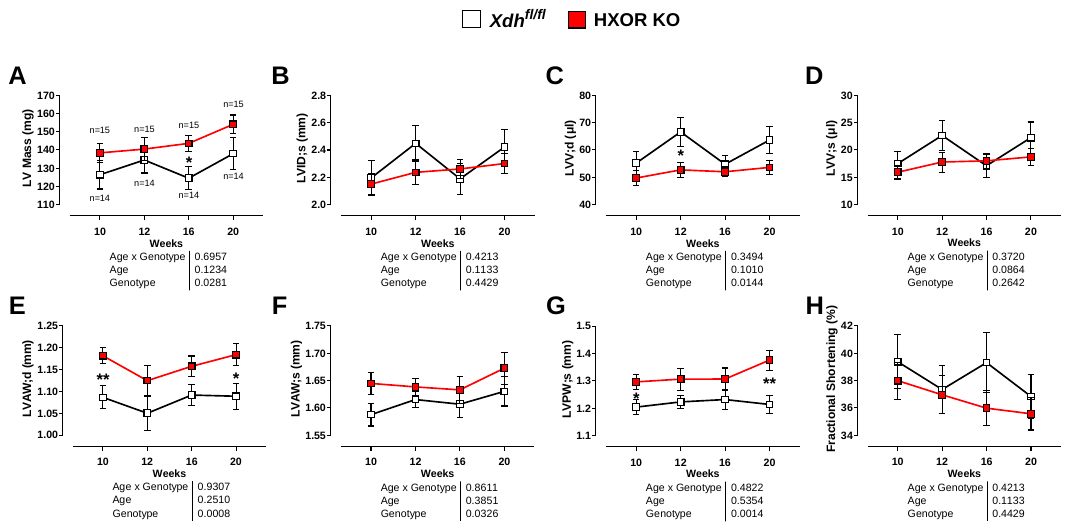
**Figure S5** HXOR KO mice have LV all thickening. Echocardiography data of *Xdh^fl/fl^* versus HXOR KO littermates, at 10-20 weeks old. Images were acquired from the LV of mice in short axis view with M-mode measurements for: **(A)** LV Mass, (**B)** LV internal diameter: systole, **(C)** LV volume; diastole, **(D)** LV volume; systole, (**E)** LV anterior wall; diastole, **(F)** LV anterior wall; systole, **(G)** LV posterior wall; systole, and **(H)** fractional shortening. Data are shown as mean ± SEM of n mice (shown in **(A)** applies to all graphs). Statistical significance was determined using Two-way ANOVA followed by Sidak’s multiple *post hoc* tests. *Post hoc* test at an individual timepoint (*Xdh^fl/fl^ vs* HXOR KO) where relevant is shown as *P<0.05, **P<0.01.

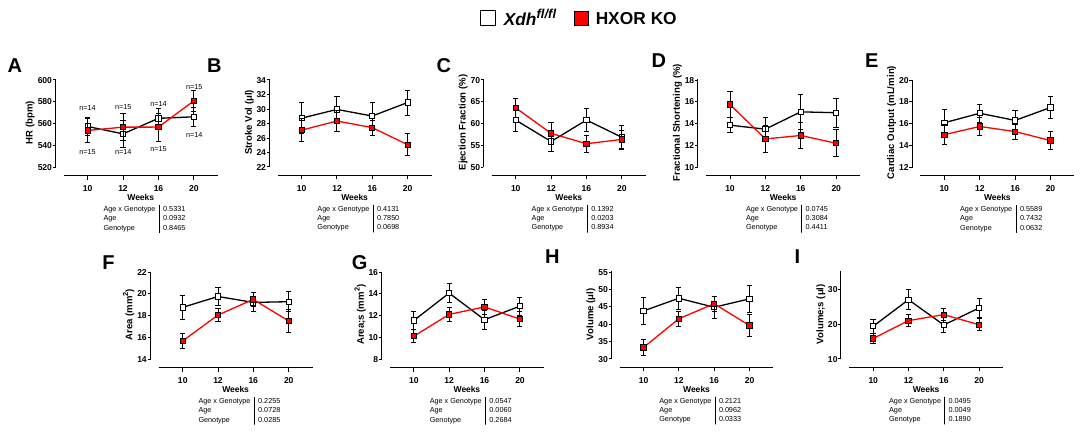
Figure S6. Confirmation of M-mode data that HXOR KO mice have LV hypertrophy and remodelling**.** Echocardiography data of *Xdh^fl/fl^* versus HXOR KO littermates, at 10-20 weeks old. Images were acquired from the LV of mice in parasternal long axis view with B-mode measurements: **(A)** heart rate, **(B)** stroke volume, **(C)** ejection fraction, **(D)** fractional shortening, **(E)** cardiac output, **(F)** LV area, **(G)** LV area: systole, **(H)** LV volume, and **(I)** LV volume; systole. Data are shown as mean ± SEM of n mice shown in **(A)** applies to all graphs. Statistical significance was determined using Two-way ANOVA followed by Sidak’s multiple *post hoc* tests.

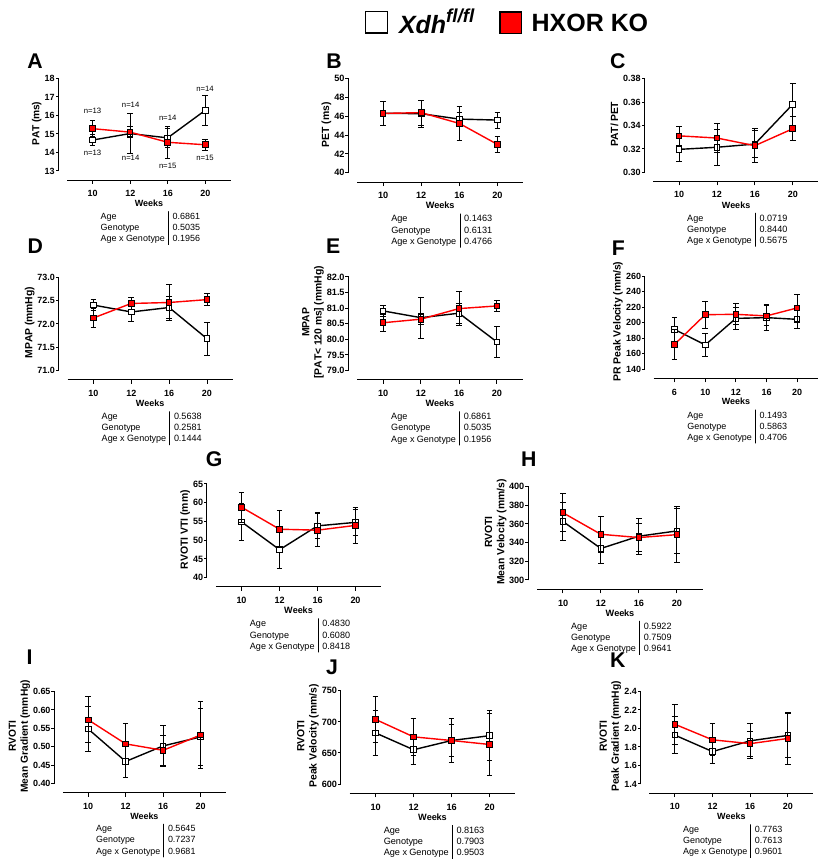
Figure S7. HXOR KO Mice have normal pulmonary artery flow**.** Echocardiography data of *Xdh^fl/fl^* versus HXOR KO littermates, at 10-20 weeks old. Values are acquired from the doppler wave measurement of pulmonary artery outflow tract for: **(A)** Pulmonary acceleration time (PAT), **(B)** pulmonary ejection time (PET), **(C)** PAT/PET, **(D)** mean pulmonary arterial pressure (mPAP), **(E)** MPAP with PAT under 120 ms, **(F)** pulmonary regurgitation peak velocity, **(G)** right ventricular outflow tract (RVOT) velocity time integral (VTI), **(H)** RVOT mean velocity, **(I)** RVOT mean gradient, **(J)** RVOT peak velocity, and **(K)** RVOT peak gradient. Data are shown as mean ±SEM of n mice (shown in **(A)** applies to all graphs). Statistical significance was determined using Two-way ANOVA followed by Sidak’s multiple *post hoc* tests.

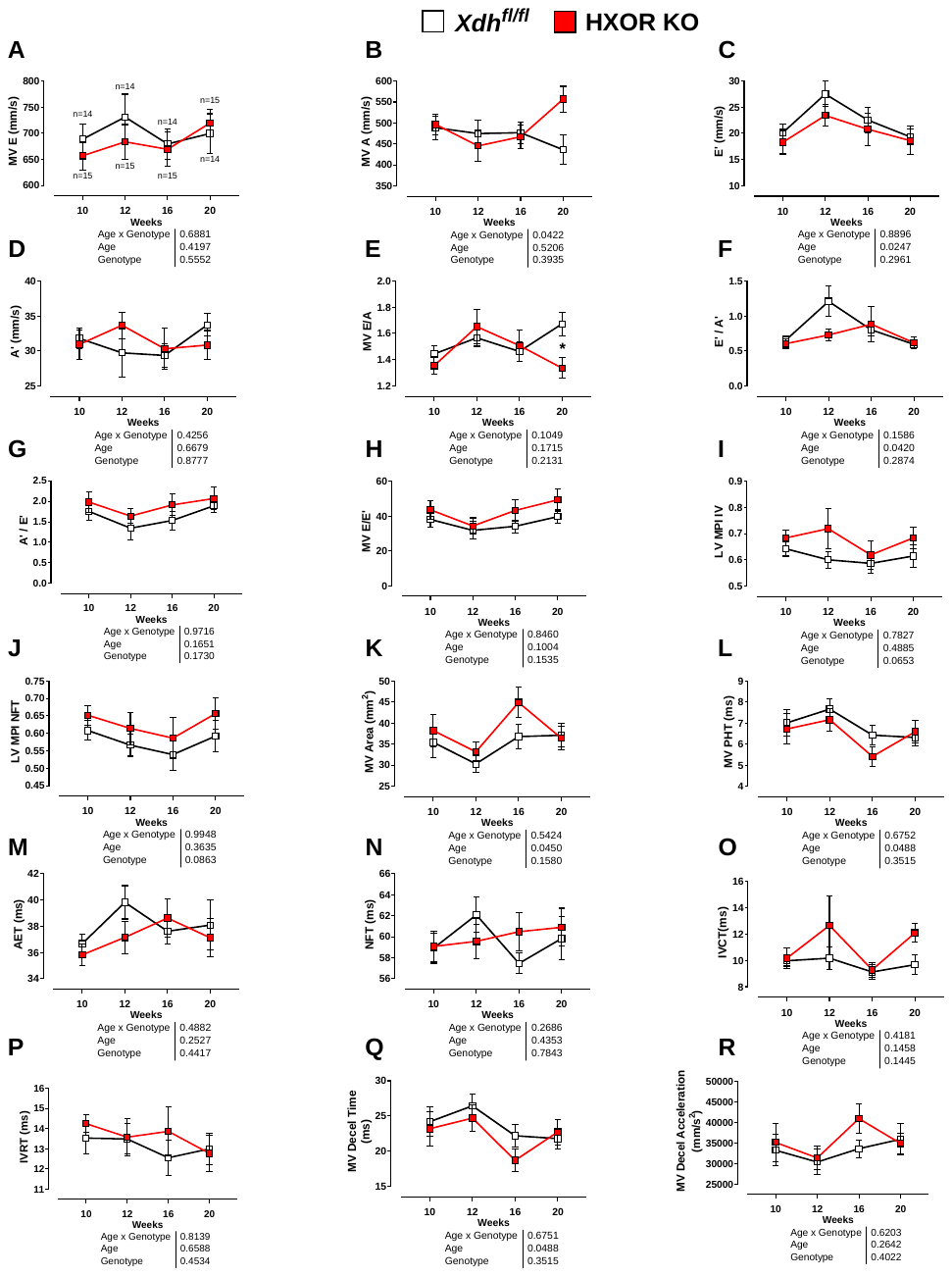
Figure S8. HXOR KO mice have a trend of impaired diastolic filling and MV flow. Echocardiography data of *Xdh^fl/fl^* versus HXOR KO littermates, at 10-20 weeks old. Data from apical 4 chamber view of the heart with tissue doppler placed on the mitral annulus and pulse wave doppler from the mitral ventricular (MV) valve were acquired; **(A)** MV E **(B)** MV A, **(C)** E’ (‘ donates prime), **(D)** A’, **(E)** MV E/A, **(F)** MV E/A’, **(G)** A’/E’, **(H)** MV E/E’, **(I)** LV myocardial performance index (MPI), **(J)** MPI No-Flow Time (NFT), **(K)** MV area, **(L)** MV Pressure Half-Time (PHT), **(M)** Aortic Ejection Time (AET), **(N)** NFT, **(O)** isovolumic contraction time (IVCT), **(P)** isovolumic relaxation time (IVRT), **(Q)** MV Deceleration time, and **(R)** MV Deceleration time rate. Data are shown as mean ±SEM of n mice (shown in **(A)** applies to all graphs). Statistical significance was determined using Two-way ANOVA followed by Sidak’s multiple *post hoc* tests.

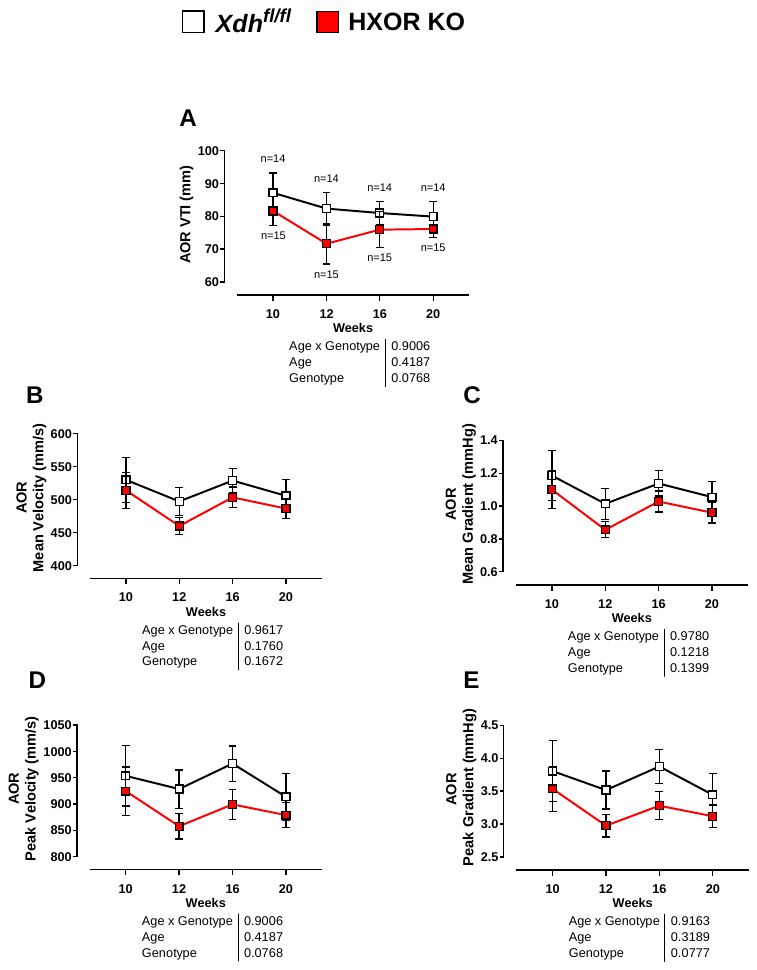

Figure S9. Trends for diastolic impairment, decreased stroke volume, and increased hypertensive afterload - corresponds with attenuated aorta VTI. Echocardiography data of *Xdh^fl/fl^* versus HXOR KO littermates, at 10-20 weeks old. Values are acquired from the doppler wave measurement of aorta outflow tract for: (A) Aorta VTI, (B) aorta mean velocity, (C) aorta mean gradient, (D) aorta peak velocity, and (E) aorta peak gradient. Data are shown as mean ±SEM of n mice (shown in (A) applies to all graphs). Statistical significance was determined using Two-way ANOVA followed by Sidak’s multiple *post hoc* tests.

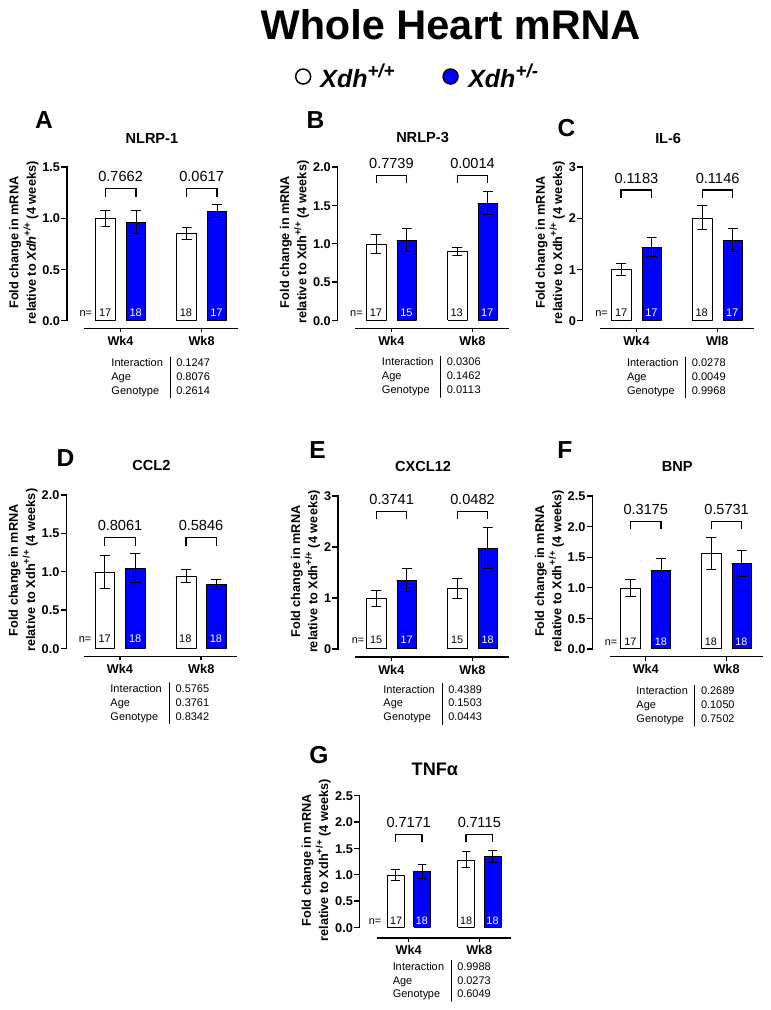

#### **Figure S10 - Cardiac mRNA expression in the Global KO model**. Longitudinal mRNA expression profiling in total mRNA extracted from whole hearts of *Xdh^+/+^* versus *Xdh^+/-^* mice at 4 and 8 weeks old investigating markers of inflammation; (A) NLRP-1, (B) NLRP-3, (C) IL-6, (D) CCL2, (E) CXCl12, (F) BNP, (G) and TNFα. Data are shown as mean ± SEM of n mice (shown in each individual graph). Statistical significance was determined using mixed-effect analysis followed by Sidak’s multiple *post hoc* tests. Uneven n values relate to technical failures or exclusion using ROUTS test.

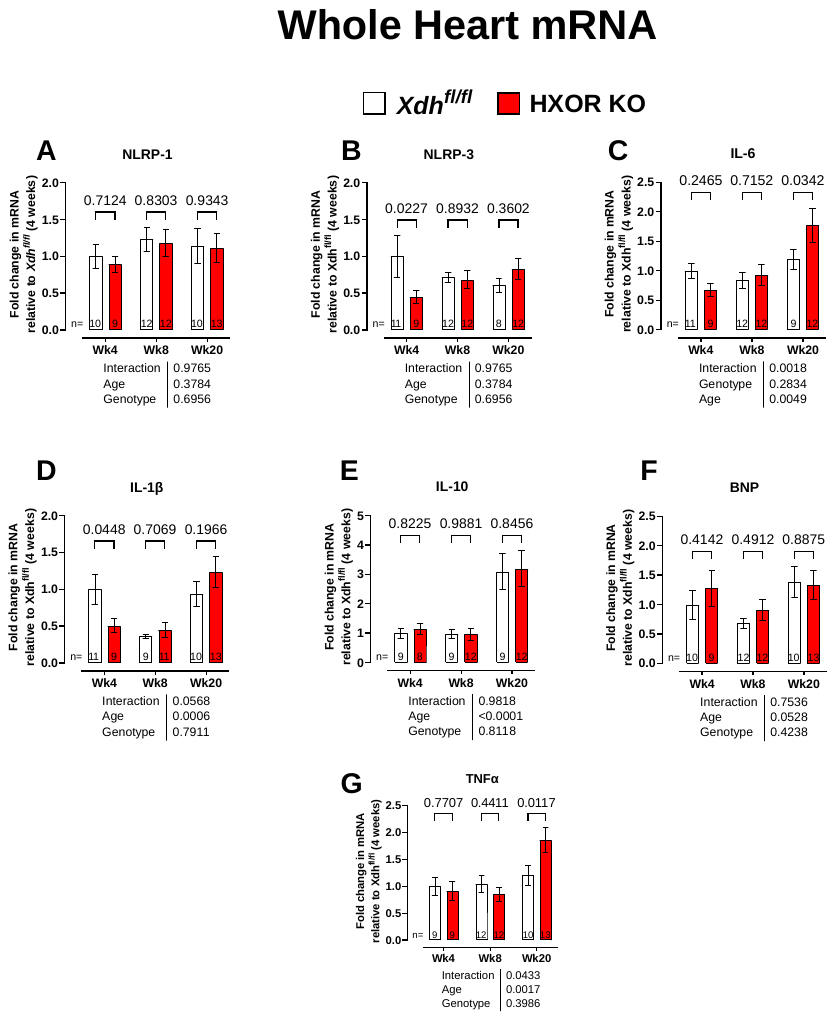
Figure S11 - Increased expression of cardiac proinflammatory mRNA in HXOR KO model**.** Longitudinal mRNA expression profiling in total mRNA extracted from whole hearts of *Xdh^fl/fl^* versus HXOR KO mice at 4, 8, and 20 weeks old investigating markers of inflammation; **(A)** NLRP-1, **(B)** NLRP-3, **(C)** IL-6, **(D)** IL-1β, **(E)** IL-10, **(F)** BNP, and **(G)** TNFα. Data are shown as mean ± SEM of n mice (shown in each individual graph). Statistical significance was determined using mixed-effect analysis followed by Sidak’s multiple *post hoc* tests. Uneven n values relate to technical failure or exclusions using ROUTS test.

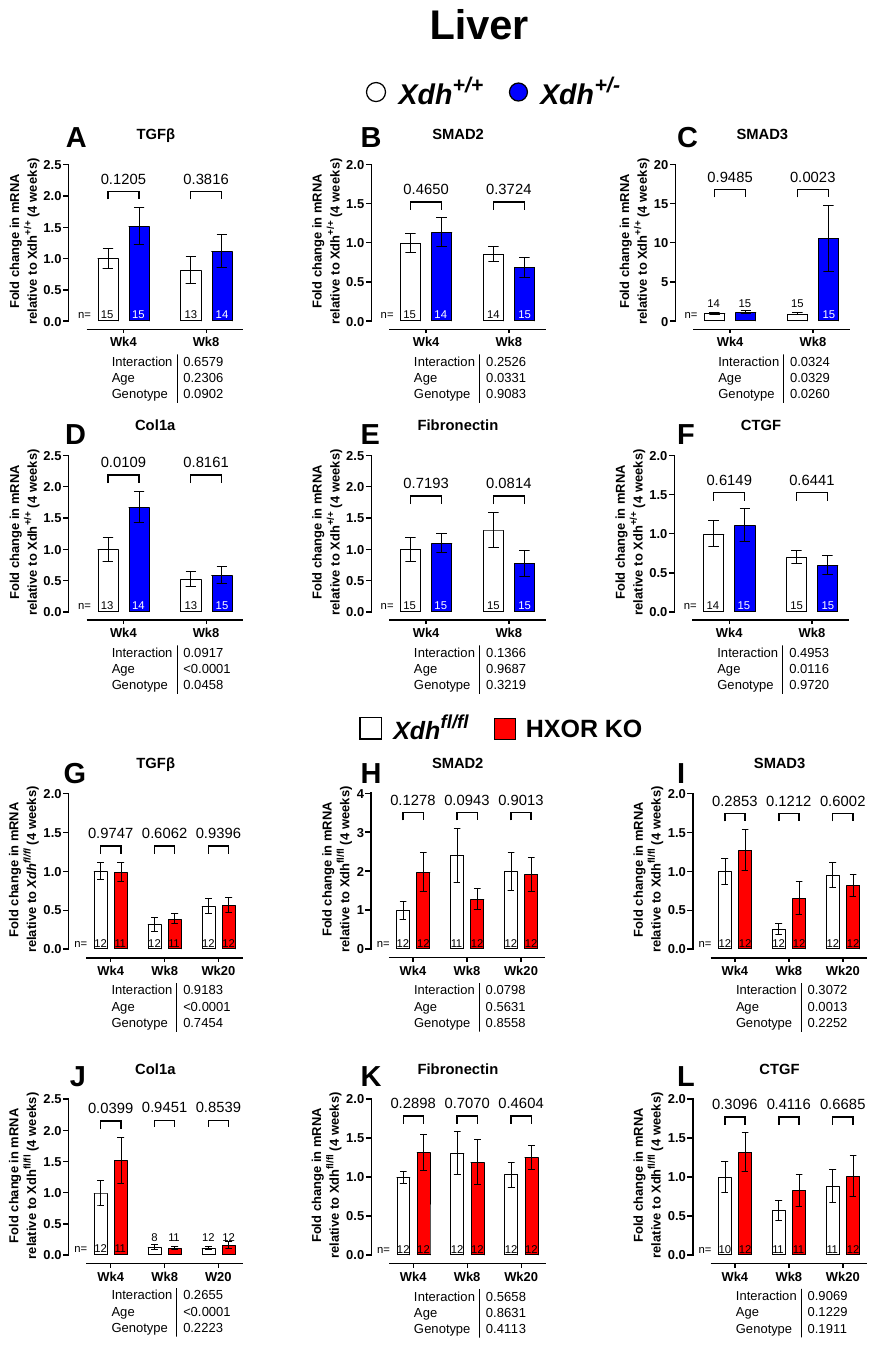

Figure S12 - Genetic deletion of hepatic XOR increase pro-fibrotic signalling pathways. Longitudinal mRNA expression profiling in total mRNA extracted from the livers of *Xdh^+/+^* versus *Xdh^+/-^*, and *Xdh^fl/fl^* versus HXOR KO mice at 4, 8, and 20 weeks old investigating markers of fibrosis; **(A,G)** TGF-β, **(B,H)** SMAD2, **(C,I)** SMAD3, **(D,J)** Col1a, **(E,K)** Fibronectin, and **(F,L)** CTFG. Data are shown as mean ±SEM of n mice (shown in each individual graph). Statistical significance was determined using mixed-effect analysis followed by Sidak’s multiple *post hoc* tests. Uneven n values relate to technical failure or exclusions via ROUTS test.

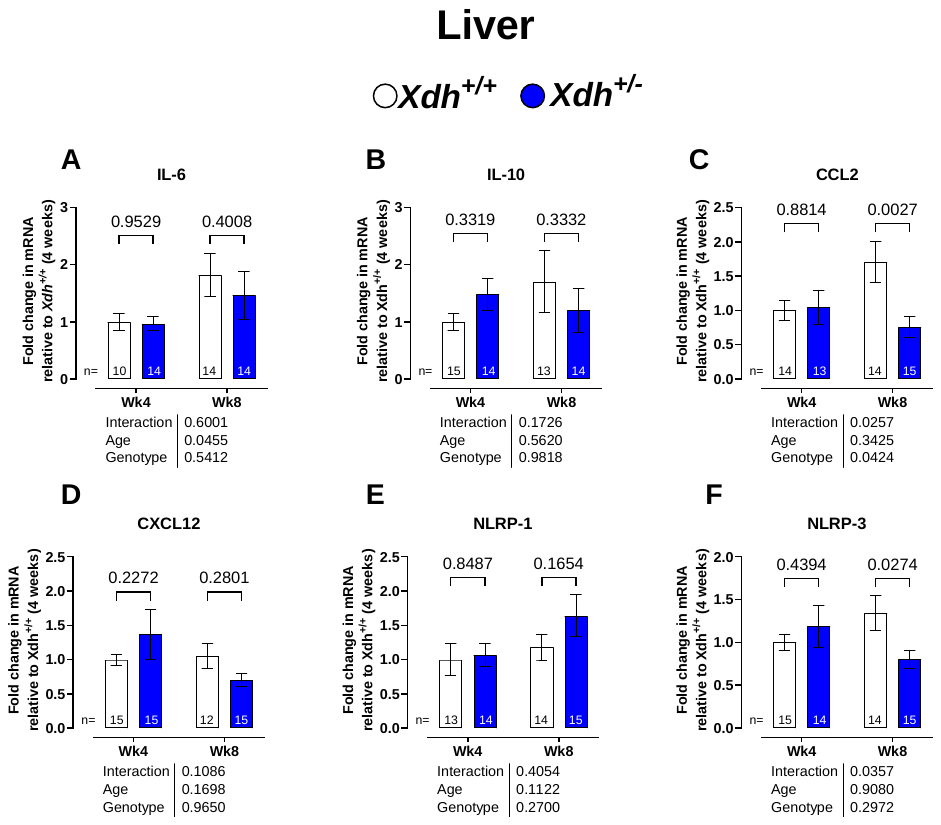
**Figure S13 – No change in mRNA markers of liver fibrosis and inflammation in Xdh^+/-^ mice.** Longitudinal mRNA expression profiling in total mRNA extracted from the livers of *Xdh^+/+^* versus *Xdh^+/-^* mice at 4-, and 8-weeks old investigating markers of inflammation; **(A)** IL-6, **(B)** IL-10, **(C)** CCL2, **(D)** CXCL12, **(E)** NLRP1, and **(F)** NLRP3. Data are shown as mean ± SEM of n mice (shown in each individual graph). Statistical significance was determined using mixed-effect analysis followed by Sidak’s multiple *post hoc* tests. Uneven n values relate to technical failure or exclusions via ROUTS test.

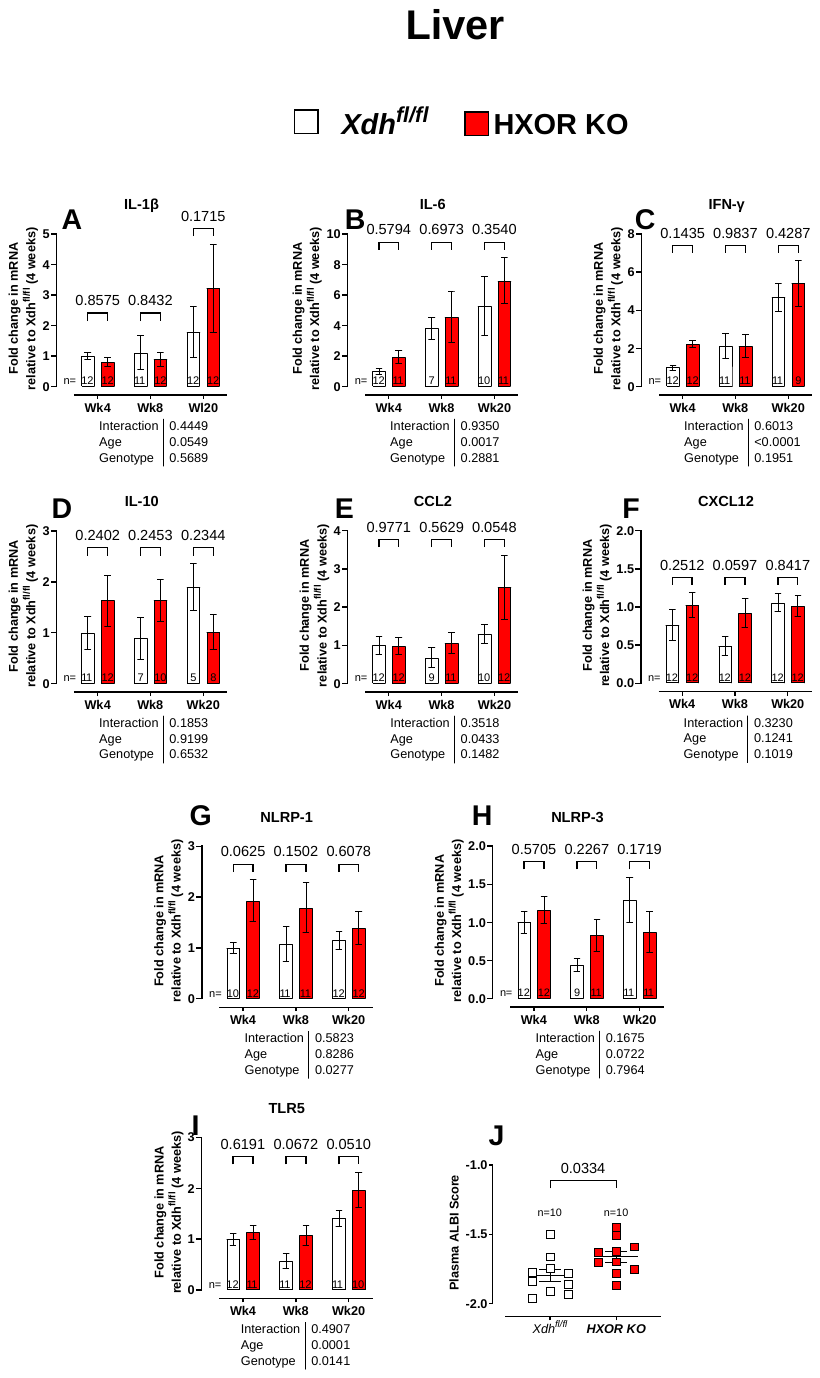
**Figure S14 – No change in mRNA markers of liver fibrosis and inflammation in HXOR KO mice.** Longitudinal mRNA expression profiling in total mRNA extracted from the livers of *Xdh^fl/fl^* versus HXOR KO mice at 4-, 8-, and 20- weeks old, investigating markers of inflammation; **(A)** IL-1β, **(B)** IL-6, **(C)** IFN-γ, **(D)** IL-10, **(E)** CCL2, **(F)** CXCL12, **(G)** NLRP1, **(H)** NLRP3, and **(I)** TLR5. **(J)** Albumin-bilirubin (ALBI) score = (−0.085 × (albumin g/L) + 0.66 × l g (TBil μmol/L)) from the values collected in plasma of 20-week-old HXOR KO mice (Table S2). Data are shown as mean ± SEM of n mice (shown in each individual graph). Statistical significance was determined using mixed-effect analysis followed by Sidak’s multiple *post hoc* tests**(A-I) or** using unpaired Student’s t-test **(J)**. Uneven n values relate to technical failure or exclusions via ROUTS test.

**Figure S15 – No change in hepatic CD45+ expressing cells in *Xdh^+/-^* and HXOR KO mice**. **(A)** Representative liver sections and histological staining of; CD45+ cells, perivascular and interstitial collagen via PSR staining of 8-week-old *Xdh^+/+^* versus *Xdh^+/-^*, and 8-, and 20-week-old *Xdh^fl/fl^* versus HXOR KO mice. Scale bar = 50 μm for CD45+ and PSR staining. **(B)** Quantification of histological analysis of 8-week-old *Xdh^+/-^* CD45+ cells. **(C)** Quantification of histological analysis of PSR perivascular collagen and **(D)** PSR interstitial collagen. **(E)** Quantification of histological analysis of 8-week-old and 20-week-old HXOR KO liver for CD45+ cells. **(F)** Quantification of histological analysis of PSR perivascular collagen and **(G)** PSR interstitial collagen. Data are shown as mean ± SEM of n mice (shown in each individual graph). Statistical significance was determined using unpaired Student’s t-test **(B-D)** or using mixed-effect analysis followed by Sidak’s multiple *post hoc* tests **(E-G).**

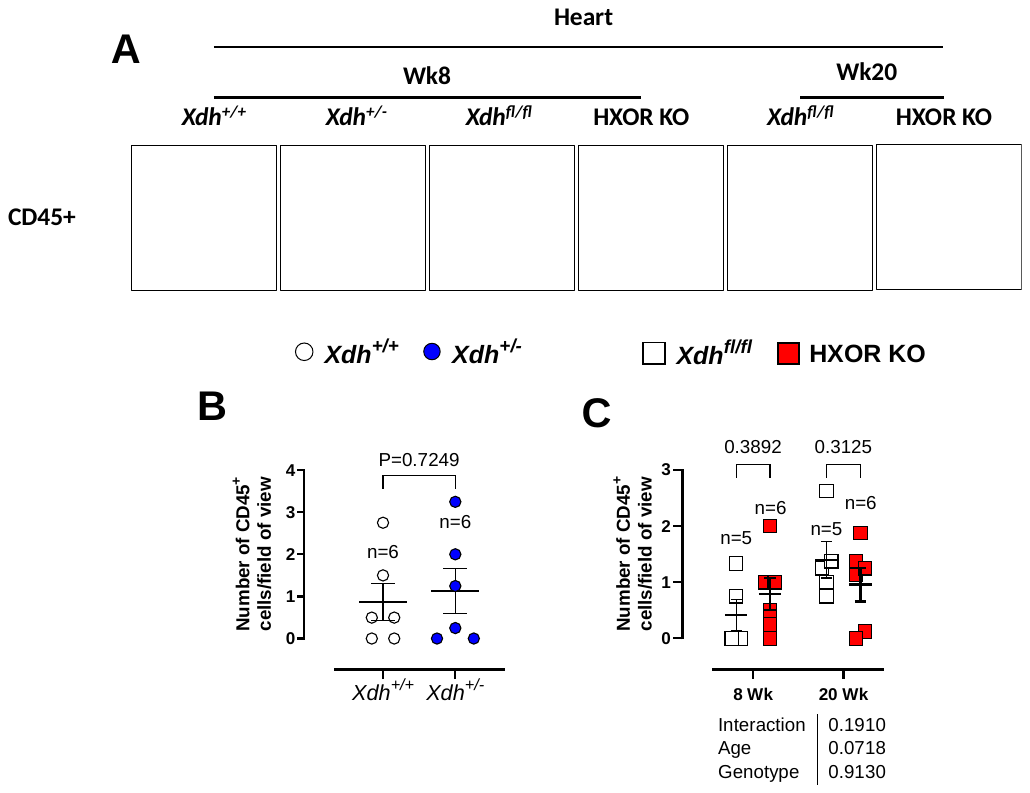
**Figure S16 – No change in cardiac CD45+ expressing cells in *Xdh^+/-^* and HXOR KO mice**. **(A)** Representative LV sections and histological staining of CD45+ staining in; 8-week-old *Xdh^+/+^* versus *Xdh^+/-^*, and 8-, and 20-week-old *Xdh^fl/fl^* versus HXOR KO mice. Scale bar = 50 μm for CD45+ staining. **(B)** Quantification of histological analysis of CD45+ cells in 8-week-old *Xdh^+/-^* mice and in **(C)** 8 and 20-week-old HXOR KO mice **(B)** Statistical significance was determined using unpaired Student’s t-test or **(C)** Two-way ANOVA followed by Sidak’s *post hoc* analysis versus *Xdh*^fl/fl^ at respective timepoint.

### References

#1.	Gu Z, Eils R, Schlesner M. Complex heatmaps reveal patterns and correlations in multidimensional genomic data. Bioinformatics (Oxford, England) 2016;**32**(18):2847-9.

#2.	Krämer A, Green J, Pollard J, Jr., Tugendreich S. Causal analysis approaches in Ingenuity Pathway Analysis. Bioinformatics (Oxford, England) 2014;**30**(4):523-30.
